# Troxerutin acts on complement mediated inflammation to ameliorate arthritic symptoms in rats

**DOI:** 10.1101/2020.08.18.253427

**Authors:** Debasis Sahu, Subasa Chandra Bishwal, Md. Zubbair Malik, Sukanya Sahu, Sandeep Rai Kaushik, Shikha Sharma, Ekta Saini, Rakesh Arya, Archana Rastogi, Sandeep Sharma, Shanta Sen, R. K. Brojen Singh, Ranjan Kumar Nanda, Amulya Kumar Panda

## Abstract

Troxerutin (TXR), is a phytochemical reported to possess anti-inflammatory and hepatoprotective effects. In this study, we aimed to exploit anti-arthritic properties of TXR using an adjuvant induced arthritic (AIA) rat model. AIA induced rats showed highest arthritis score at disease onset and by oral administration of TXR (50, 100, 200 mg/kg body weight), reduced to basal level in a dose dependent manner. Isobaric tag for relative and absolute quantitative (iTRAQ) proteomics tool was employed to identify deregulated joint homogenate proteins in AIA and TXR treated rats to decipher probable mechanism of the TXR action in arthritis. iTRAQ analysis identified a set of 434 joint homogenate proteins with 65 deregulated proteins (log_2_ case/control ≥1.5) in AIA. Expressions of a set of important proteins (AAT, T-kininogen, vimentin, desmin, and nucleophosmin) that could classify AIA from healthy were validated using Western blot analysis. Western blot data corroborated proteomics findings. *In silico* protein-protein interaction study of joint homogenate proteome revealed that complement component 9, the major building blocks of the membrane attack complex (MAC) responsible for sterile inflammation, gets perturbed in AIA. Our dosimetry study suggests that a TXR dose of 200 mg/kg body weight for 15 days is sufficient to bring the arthritis score to basal levels in AIA rats. We have shown the importance of TXR as an anti-arthritis agent in AIA model and after additional investigation its arthritis ameliorating properties could be exploited for clinical usability.

## 1. Introduction

Rheumatoid arthritis (RA) is a chronic, systemic autoimmune inflammatory disorder primarily affecting the synovial joints with concomitant destruction of joint tissues. Almost 1% of the world population irrespective of race and region is suffering from RA with its preponderance more in females than males^1^. Commonly used anti-tumor necrosis factor (TNF) drugs provide relief to only about 60% of the RA patients. Long term use of nonsteroidal anti-inflammatory drugs (NSAIDs) such as diclofenac sodium to manage RA leads to persistent adverse events^2^. Therapeutic advances have significantly improved the lives of RA patients but the problem needs additional solutions to resolve the clinical issues associated with this disease. Identification of alternate drugs with minimal toxicity is the need of the hour. Plant based extracts and phytochemicals have been used as a rich source of anti-RA agents^3–5^.

Troxerutin (TXR), also known as vitamin P4, is a flavonoid found in cereal, coffee, tea, many vegetables/fruits and reported to have a wide range of pharmacological properties^6,7^. TXR protects oxidative damage of cell membranes of neutrophils, DNA damage caused by gamma radiations, protects renal and hepatic conditions. TXR reduces capillary fragility and unusual leakage thus improving their function. It is reported to possess fibrinolytic, anti-thrombotic, rheological and edema-protective activity^8^. TXR has also been used to treat chronic venous insufficiency (CVI)^9,10^.

In this study, we monitored the antioxidant potential of TXR in an in-vitro system and to decipher its arthritis ameliorating potential, an adjuvant induced arthritis (AIA) model of rat was used for monitoring morphological, histological, radiological and biochemical parameters. A global quantitative proteomics study was employed to identify the deregulated proteins in affected synovial joint homogenate isolated from AIA induced rats and those receiving TXR treatment and validated using Western blot analysis. The identified joint homogenate proteins were mapped to identify interacting partners and used to predict the mechanism of TXR action. This study successfully demonstrated the importance of phytochemicals like TXR as anti-arthritis agent and elucidated its mode of action to develop translatable solutions for difficult debilitating disease conditions like RA.

## 2. Methodology

### 2.1. Cell culture

Murine macrophage cell-line (RAW264.7) was cultured using RPMI 1640 (Gibco, USA) medium supplemented with glutamine (2 mM), antibiotics (streptomycin: 100 μg/ml, amphotericin-B: 0.25 μg/ml and penicillin: 100 U/ml), and fetal bovine serum (heat-inactivated, 10%) in a humidified CO_2_ incubator maintained at 37°C. Varied TXR concentrations (5.2 to 674 μM) were introduced to RAW264.7 cells with controls, for 24 hours. Trypan blue (0.4% in phosphate buffered saline: PBS) exclusion method was used to determine cell viability in TXR treated cells. Cytotoxicity was measured using MTT colorimetric assay (Sigma Aldrich, USA), and Griess-nitrite assay (Sigma Aldrich, USA) was carried out to measure nitric oxide production with sodium nitroprusside (SNP, 75μM, Sigma Aldrich, USA) as reported earlier with minor modifications^11^.

### 2.2. Animal experiments

All animal experiments were performed following approved protocols by the Institutional Animal Ethics Committee (IAEC) of National Institute of Immunology (NII), New Delhi (IAEC#367/15). Female Wistar rats, aged 6-8 weeks were maintained at controlled temperature (24±3°C) and humidity (50±5%) with 12 h light and dark cycles; and had access to food and water *ad libitum*. After a period of acclimatization (7 days) in the animal house, study animals were randomly divided into six groups (n=5/group). Schematic diagram depicting the experimental procedures adopted in this study is presented in Figure 1 (1A, 1B, 1C and 1D). Except the healthy control groups, rest five groups were immunized with complete Freund’s adjuvant (CFA; 100 μl, Chondrex, USA) by intradermal injection in the subplantar region of both hind footpads. Post CFA administration, animals were monitored for 24-36 hours and the footpad thickness was recorded using digital Vernier calipers (Mitutoyo, Japan). These animals were divided into the following groups (vehicle control, AIA, diclofenac sodium control (1 mg/kg DS) and treated with different doses of TXR (50 mg/kg: TXR 50; 100 mg/kg: TXR 100 and 200 mg/kg: TXR 200)) administered orally once a day for 15 days. Body weights of experimental animals were recorded every 3^rd^ day post AIA induction till euthanization and the difference in mean body weight at two representative days (5 and 21) were calculated. Using retro-orbital puncture, blood samples (~0.5 ml) were collected on day 20. After euthanizing the animals, their hind limbs were harvested for x-ray imaging, histology and proteomic studies. Harvested kidneys and livers fixed in formalin (10% v/v in phosphate buffered saline: PBS) for histopathological analyses. For radiological evaluation, right hind-limbs of each experimental animal from all study groups were fixed in formalin (10% v/v in PBS). Joint tissues excised from the left hind limbs of the animals were snap-frozen to store at −80°C for further analyses.

**Figure 1:**
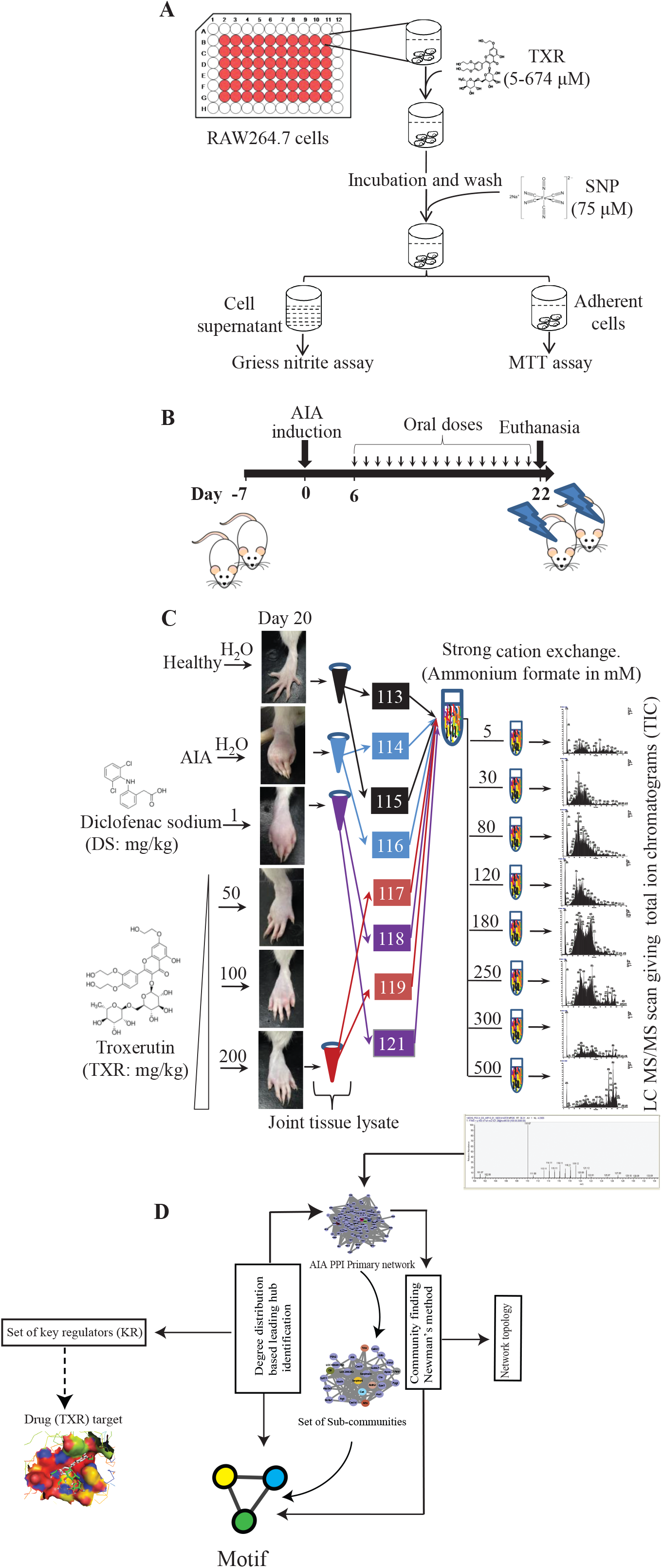
Schematic representation of the experimental approach used in this study. A. Nitric oxide inhibition assay simultaneously with the cytotoxicity of TXR was evaluated using RAW264.7 cells. B. Induction of adjuvant induced arthritis (AIA) and the treatment plan. C. Joint homogenates from multiple study groups were used for proteomics experiment. D. Informatics analyses were used for identifying the key regulator proteins.

Prior to histology, antero-posterior roentgenograms were recorded on x-ray films using MBR-1505R (Hitachi Medical Corporation, Tokyo, Japan) at 30kV, 6 mA and 45 seconds exposure with a 50 cm distance between X-ray source and film. The severity of bone erosion was ranked according to Larsen scoring method with minor modifications. A score zero (0) was assigned to normal joints and bones, while a slight abnormality in any one or both exterior metatarsal bones exhibiting minor bone erosion was scored as 1. An image with distinct abnormality in any of the metatarsal or tarsal joints having bone erosion was scored 2. Score 3 indicated medium destructive abnormalities in the metatarsal bones or tarsal bones (one or both) showing definite bone erosions. When a severe destructive abnormality in all the metatarsal bones showing definite erosion involving at least one of the tarsometatarsal joints entirely eroded, but some bony joint outlines were partly preserved and scored as 4. The highest score was 5, assigned to severely damaged states with mutilating abnormality with no bony outlines^12^.

After the radiological assay, the knee joint was separated out from the formalin fixed limb by removing the skin and overlying muscles for histological analysis. Decalcification was carried out in nitric acid (5%) incubated for 10 days prior to paraffin block preparation. Tissue sections (5 μm) were prepared using a microtome (Leica Biosystems, Germany), stained with hematoxylin and eosin (H-E) before capturing images using light microscope (Nikon, USA)^13^.

### 2.3. Histological analysis of liver, kidney and tibiotarsal joints

Fixed liver and kidney tissues were sectioned (8 μm) using a rotary microtome (Leica Biosystems, Germany) and stained with H-E staining for visualization. For the histological analysis of liver, the following parameters were considered for the eosinophilic infiltration assay. The scoring was done according to the eosinophils counted per frame of the photomicrograph. Score 0 was given if the eosinophils were absent, with 1-2 eosinophils were scored as 1. Score 2 was assigned for ≥3 but <6 eosinophil count and the highest score of 3 was assigned for eosinophilic count ≥6. Semiquantitative scores (0-3) represent an average of 3 highest values of eosinophils, counted in at least 8 portal / periportal spaces. The histological scoring for the tibiotarsal joints and other soft tissues (liver and kidneys) was performed by a clinical pathologist using a standardized method^14,15^.

### 2.4. Measurement of nitrite in plasma

Blood (~0.5 ml) was drawn from rats by cardiac puncture immediately after euthanasia and collected in EDTA-coated vacutainers (Greiner Bio-One) for plasma nitrite estimation. Blood plasma was separated by centrifugation at 2,000 *g* for 10 min at 4°C before storing at −80°C until further analysis. Briefly, plasma (0.1 ml) and varied concentrations of sodium nitrite (100 - 1.562 μM) were taken in 96-well microtitre plate and Griess reagent (0.1ml, Sigma Inc., USA) was added to the mixture. The reaction mixture was incubated for 10 minutes in the dark before measuring the absorbance at 540 nm using a spectrophotometer (Molecular devices, USA). All samples were measured in duplicates to quantify the nitrite levels^16^.

### 2.5. Assessment of arthritis score at disease onset and during treatment

Every third day, arthritic score of all the experimental animals was recorded by a coauthor blinded to the study groups. Each experimental rat was given a score in the range of 0-4; where ‘0’ shows no symptoms and ‘4’ being the most severe. Combined score of both the hind limbs were used for analysis. The paw diameters were measured every 3^rd^ day from the date of AIA induction (day 0) to the date of euthanization (day 21) to calculate the arthritis index (AI) using equation 1^17^.

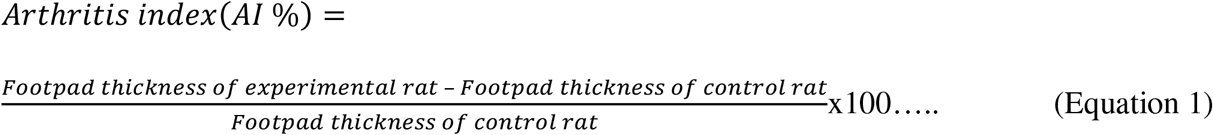

### 2.6. Proteome analysis of arthritic tissues

After clearing the skin and extra muscles, in presence of liquid nitrogen, joint tissues were pulverized to powder form using a pestle and mortar. The homogenized joint tissue samples (~500 mg each) were suspended in protein extraction buffer (50 mM Tris-HCl, pH 7.4, 1 mM PMSF and protease inhibitor cocktail; Sigma, USA Cat# P8340) for 30 min in ice before centrifuging at 4°C at 10,000 *g* for 10 min. The supernatant was used for estimating protein following Bradford’s method (5000006, Bio-Rad, USA). Equal amount of joint homogenate protein (100 μg) from each study group was precipitated in acetone (1/6 v/v) for overnight and centrifuged at 12,000 *g* for 20 min at 4°C to collect the pellet. The protein pellet was dissolved in dissolution buffer and denaturant supplied with the isobaric tag for relative and absolute quantification (iTRAQ) reagent kit (AB Sciex, USA) (Fig S1A). Equal amount of proteins from all groups were separated on a gel and silver stained gel image is presented in Fig S1B. Supplier’s protocol was followed for iTRAQ labeling. Briefly, reducing reagent (2 μl) was added to each protein sample (100 μg) and incubated at 60°C for 1 h followed by the addition of cystine blocking reagent (1 μl). Tryptic peptides were generated by incubating the processed proteins with trypsin (20 μl) followed by overnight incubation at 37°C and dried using a vacuum evaporator (CentriVap concentrator, Labconco, USA) at 40°C for 35 min. The tryptic peptides were resuspended in dissolution buffer and appropriate iTRAQ tags were used for labeling the tryptic peptides by incubating for 2 hours at room temperature (Fig. S1). Pooled iTRAQ labeled peptides were dried using SpeedVac at 40°C and an aliquot (~300 μg tryptic iTRAQ labeled peptides) was taken for strong cation exchange chromatography (SCX). Briefly, labeled peptides were dissolved in 2 ml of SCX buffer A (5 mM ammonium formate in 30% acetonitrile, pH 2.7) and loaded onto a pre-equilibrated SCX column ICAT™ cartridge kit (AB Sciex, USA). Peptides were eluted into 8 fractions using ammonium formate buffer (pH 2.7) of different concentrations (30, 80, 120, 180, 250, 300, 400, and 500 mM). Fractions derived from the 30 mM and 80 mM concentrations were combined and the rest 6 peptide fractions were separately dried in the CentriVap concentrator at 40°C. These fractions were subjected to cleaning up using Pierce C18 Spin Columns (89870, Thermo Fisher, USA).

**LC MS/MS data acquisition:** Each cleaned up fraction was taken for reverse phase separation coupled to an online mass spectrometer for data acquisition. Briefly, peptide fractions (~ 20 μg) were dissolved in 20 μl solvent A (5% acetonitrile with 0.1% formic acid) of which 2 μl (2 μg) was injected into a precolumn and separated on a 10 cm C_18_ Pico Frit analytical column, Hypersil Gold 5μ, 15 micron tip with an internal ID of 75 micron (PF7515-100H-070-36, New Objective, USA). In a nano-LC-MS column (Thermo Fisher Scientific, USA), peptides were separated via a constant flow rate of 300 nl/min with a mixture of solvent A and solvent B (95% acetonitrile with 0.1% formic acid) to achieve a solvent gradient of 5 to 38% acetonitrile in 70 min, then up to 76% acetonitrile from 70 to 80 min, and maintained until the completion of the 120 min run time. Mass spectrometer data acquisition was carried out using a Linear Trap Quadrupole (LTQ) Orbitrap Velos mass spectrometer (Thermo Fisher Scientific, USA). The eluates of nano-LC were directly sampled via an integrated electrospray emitter operating at 2.0 kV. Full-scan (m/z: 200–2000 MS spectra) data was acquired in the positive ion mode with a ten (10) data-dependent collision-induced dissociation (CID) - higher-energy collisional dissociation (HCD) dual MS / MS scans per full scan using Fourier transform mass spectrometry (FTMS) mass analyzer; CID scans were obtained in LTQ with two-microscan averaging; HCD scans and full scans were acquired in the Orbitrap at a resolution of 15,000 and 60,000 respectively; normalized collision energy (NCE) was of 50% in HCD and 30% in CID; ±2.0 *m*/*z* isolation window; and the dynamic exclusion for 60s. In CID-HCD dual scan, selected parent ions were fragmented by CID followed by HCD. The peptides with charged states of +2 or more were considered for MS/MS fragmentation. The 10 major high abundant peptides, with 500 and above signal threshold count were selected for MS/MS and excluded dynamically for 30 milliseconds. Protein identification and quantification: All the mass spectrometric raw data files were analyzed using SEQUEST or MASCOT search algorithm and peak list generation in Thermo Proteome Discoverer 1.3.0.339 software (Thermo Fisher Scientific, USA). Thermo Xcalibur Qual Browser was used as the search engine with Uniprot-rat.fasta (Database: Uniprot; Species: *Rattus norvegicus* (Rat); Taxon identifier: 10116; as accessed on 14-15/03/2018) as the sequence database. Following search parameters like mass tolerance for precursor and fragment ion at 10 ppm and 0.1 Da respectively were selected for analysis. Enzyme specificity was set to trypsin with less than two missed cleavage sites, static modification (peptide N-terminus) with iTRAQ 8 plex / +304.205 Da (K); methylthio / 45.988 Da (C), dynamic modification with +15.995 Da (M) (oxidation). All identifications were filtered using the peak integration window tolerance of 20 ppm and analysis of the top 10 peaks. Estimation of false discovery rate (FDR) was calculated using the parameter of target FDR as 0.01 at peptide level. Proteins with at least two peptides were uniquely assigned to the respective sequence and were considered for further analysis at 95% significance level. All qualified proteins were exported to Microsoft Excel for manual data interpretation. Fold change of proteins were presented in logarithmic scale and proteins with ≥1.5 fold difference (log_2_ fold change ≥ ±0.585) and *p* ≤ 0.05 were considered as important deregulated proteins and selected for further analysis. A minimum set of selected proteins as important molecular signature were selected for validation using Western blot assay and monitored in the treated subjects.

### 2.7. Data availability

All raw data files were submitted in ProteomeXchange database and could be assessed using identifiers with accession numbers: PXD004982 (ProteomeXchange) or MSV000080172 (MassIVE).

### 2.8. Western blot

Equal amount of rat joint protein extracts (20 μg), from the animals not used in the discovery set, were subjected to electrophoresis on 12 % sodium dodecyl sulfate polyacrylamide gel (SDS-PAGE). Separated proteins were transferred to PVDF membrane (Amersham Biosciences, UK) at 55 mA for 1.5 hours in 25 mM Tris, 192 mM glycine, 20% methanol using a TE 77-semidry transfer unit (GE healthcare Bio-sciences, USA). Bovine serum albumin (BSA, 1% in Tris buffered saline with 0.1% Tween-20 detergent: TBST) was used for blocking the non-specific binding sites at 4°C by gentle shaking overnight. After washing with TBST buffer, protein transferred PVDF membranes were probed with respective primary antibodies (at 1:1000 dilution in TBST containing 0.25% BSA) against rat desmin (sc-23879, Santa Cruz, USA), vimentin (sc-32322, Santa Cruz, USA), T-kininogen (sc-103886, Santa Cruz, USA), alpha-1-antitrypsin (ab166610, Abcam, UK), nucleophosmin (ab10530, Abcam, UK) and GAPDH (sc365062, Santa Cruz, USA) for 3 hours at room temperature. After washing, the blots were incubated with anti-rabbit (sc-2357, Santa Cruz, USA) or anti-goat IgG secondary antibodies (sc-2354, Santa Cruz, USA) conjugated with horseradish peroxidase (1:5000 in TBST containing 0.25% BSA) at room temperature for 2 hours. The blots were washed again, equal volumes of solution A and B luminol reagents (sc-2048, Santa Cruz, USA) were properly mixed before adding to the blot and the resulting chemiluminescent signal was captured on X-ray film (Kodak, India) in a dark room. Western blots were scanned using Image Scan and Analysis System (Alpha-Innotech Corporation, USA)^18^. The band intensities were calculated using ImageJ and taken for densitometric analysis.

### 2.9. Protein-protein interaction (PPI) network construction

The set of identified joint homogenate proteins from iTRAQ experiment was used to construct a protein-protein interaction (PPI) network using Cytoscape 3.6.0^19^ appsGeneMANIA^20^. Only the physical interaction network of arthritis-related genes was extracted from the constructed PPI. The network was curated after deletion of the isolated node(s). So, a total of 434 nodes and 13,316 edges were built as a primary network graph and denoted by *G(N,E), N* represents the set of nodes with *N* = {*ni*}; *i* = 1,2, … ‥, *N* while *E* denotes the set of edges with *E* = {*eij*}; *i, j* = 1,2,3, … ., *N*.

### 2.10. Detection of the levels of organization

The communities of arthritic PPI network were extracted using community finding method of Newman and Grivan which is the first level of network organization^21^. It was established as a result of communities’ interaction from the primary PPI network. Second organizational level was constituted by the subcommunities prepared from all communities forming the first level organization. Similarly, the succeeding levels were formed with the construction of *motifs.* Thus each smaller community possesses at least one triangular *motif* as defined by sub-graph *G(3,3)*. *Motif G(3,3)* was used as the criteria of qualification for community or subcommunity as a constituting member at certain level of organization because the triangular motif was overrepresented in PPI network and served as the controlling unit in the network^22^. So, each community belongs to the different levels of organization.

### 2.11. Topological analyses of the networks

The topological properties of the network for centralities, clustering coefficients, degree distribution, and neighborhood connectivity were analyzed using Cytoscape plugins, NetworkAnalyzer^23^ and CytoNCA^24^.

**Degree(*k*):** In the process of network analysis, the total number of links is established by a node in the network and indicated by the degree *k*. this is also used to measure the local significance of a node in the network regulation process. In the graph represented by *G = (N, E)*, *N* denotes the nodes while the edges are denoted by E. The degree of *i*^th^ node (*k*_*i*_) is expressed as 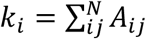, where the adjacency matrix elements of the graph is denoted by *A*_*ij*_.

**Probability of degree distribution (*P*(*k*)):** *P(k)* denotes the probability of a random node for having a degree k out of the total number of nodes present in the network. This is represented as the fraction of nodes with degree (*k*), as presented in the equation 2; here *N*_*k*_ denotes the number of nodes to have degree *k* and *N* represents the total number of nodes in the network.

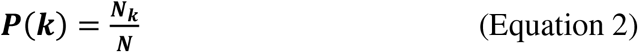

*P*(*k*) of small world and random networks always follow Poisson’s distribution in a degree distribution against degree (k), however most of the real world networks which are hierarchical and scale-free follow the power law distribution *P*(*k*) ~ *k*^*γ*^ and 4 ≥ *γ* ≥ 2. In the hierarchical networks, *γ* ~ 2.26 (mean-field value) indicates a hierarchical modular organization at different topological levels^25^. So, the characteristic topology of a network is defined by *P*(*k*) pattern.

**Clustering coefficients *C*(*k*):** It is the ratio of number of triangular motifs created by a node with its nearest neighbors and the total number of such motifs in the entire network. Therefore, *C(k)* characterizes the strength of internal connectivity within the nodes neighborhoods quantifying the inherent clustering tendencies of nodes. Equation 3 expresses *C*(*k*) for any node *i* with the degree *k*_*i*_ in a unidirectional graph. In this equation, *m*_*i*_ represents the total number of edges among its nearest neighbors. In the scale-free networks *C*(*k*)~ *constant*, however it shows the power law in a hierarchical network against the degree, *C*(*k*)~ *k*^−*α*^, with *α* ~ 1 ^25^.

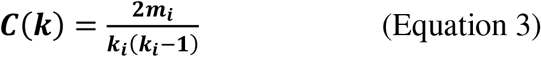

**Neighborhood connectivity *C*_*N*_(*k*):** The node neighborhood connectivity is the average connectivity which is established by nearest-neighbors of a node with degree *k*. It is represented by *C*_*N*_(*k*) as shown in equation 4, here *P*(*q*|*k*) is a conditional probability of a node’s links with *k* connections to one of the other nodes having *q* connections.

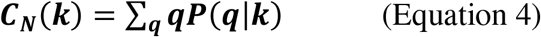

In the hierarchical network topology, *C*_*N*_(*k*) shows power law against the degree *k*, i.e., *C*_*N*_(*k*)~ *k*^*β*^, here *β*~0.5 ^26^. The negativity or positivity of the exponent *β* may be defined as disassortivity or assortivity nature of a network topology, respectively^27^.

**Centrality measures.** Betweenness centrality ***C***_***B***_, closeness centrality ***C***_***C***_, Eigenvector centrality ***C***_***E***_ are the basic centrality measures and are the parameters for the estimation for a node’s global functional significance in a network regulation through information processing ^28^.

The total geodesic distance between a node and all of its connected nodes is given by *C*_*C*_. It also determines how rapidly an information is spread within a network from one node to other connected nodes ^29^. In a given network, *C*_*C*_ of a node *i* is calculated by dividing the total number of nodes of network *n* by the summation of geodesic path lengths between the nodes *i* and *j* which is given by *d*_*ij*_ of equation 5.

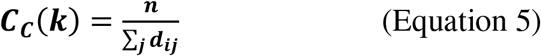

The ***C***_***B***_ or betweenness centrality is the measure of a node which is share of all the shortest-path traffic from all feasible routes through nodes *i* to *j*. So, it is the parameter of the ability of a node to extract benefit from the flow of information throughout the network^30^ and its ability to control the signal processing over the other nodes within the network ^31^. If *d*_*ij*_(*v*) represents the number of geodesic paths from one node *i* to another node *j* passing through the node *v*, then *C*_*B*_(*v*) of node *v* can be derived by the equation 6.

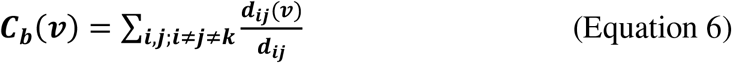

The normalized betweenness centrality is summarized in the equation 7, in which *M* represents the number of node pairs, excluding *v*.

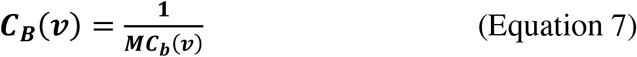

Eigenvector centrality ***C***_***E***_ corresponds to the intensity of most prominent nodes affecting signal processing throughout the network and is proportional to the sum of centralities of all neighbors of a node^32^. In a network, the nearest neighbors of node *i* is given by *nn*(*i*) with eigen value *λ* and the eigenvector *v*_*i*_ of eigen-value equations, *Av*_*i*_ = *λv*_*i*_(*v*) where, *A* is network adjacency matrix, *C*_*E*_ can be calculated by the equation 8,

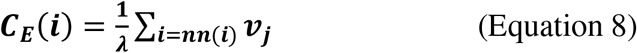

The value of *C*_*E*_ gives the maximum positive eigen value, *λ*_*max*_ of principal eigenvector of *A* ^32^. A node’s *C*_*E*_ function is dependent on the centralities of its neighbors and it varies according to different networks association of high *C*_*E*_ nodes. There are lesser chances of isolation of nodes within closely connected region of such nodes^32^. Thus, *C*_*E*_ is considered as an influential indicator of information transmission power of a node within a network.

### 2.12. Tracking the key regulators in the networks

The most prominent genes or the corresponding proteins of the arthritic network were first identified by centrality measures calculation. As the higher degree nodes possess higher centrality values, we considered the top 65 highest degree nodes (*degree k* ≥ 65) among the *hub* nodes in order to trace the key regulators which may have very important role to play in the network regulation. Then nodes were traced from primary network up to the *motif* level *G(3,3)*. This tracing was performed based on representation of respective nodes (proteins) throughout the submodules obtained from community detection / clustering method of Louvain. Therefore, the key regulators of the arthritic network were the *hub*-nodes (proteins) which corresponded to the modules at all the hierarchical levels in the arthritic network.

### 2.13. Molecular docking study

The identified key regulators were studied for their molecular interaction with troxerutin. The structures of the proteins were either taken from the Protein Data Bank (PDB) or the primary sequence of key regulators is retrieved from NCBI for homology modeling. iTASSER servers were instrumental for homology modelling of the key regulator proteins^33^. The docking study is performed using AUTODOCK vina, Discovery Studio, Schrödinger Glide software and the interactions were visualized with PYMOL^34^, Chimera^35^, Discovery Studio Visualization (Accelrys, San Diego, CA, USA), Maestro^36^.

### 2.14. Statistical analysis

All samples (cell and animals) were randomly selected for each experimental group. Based on our experience, the sample and animal sizes were selected in order to achieve sufficient statistical power to identify relevant differences, if any. Based on the distribution of data and the number of study groups used for comparative analysis, appropriate statistical tests were explored for determining the levels of significance. Two ways analysis of variance (ANOVA) with Bonferroni post-test corrections for multiple comparisons with uncorrected Fisher’s LSD test were used for determining the difference between AIA and other groups, for AIA score, arthritic index, change in footpad thickness and body weight. Whereas, one way ANOVA with Dunnett’s post tests for multiple comparisons was used for radiographic score, all the histological scores, Griess assay in cell culture, MTT assays, and western blotting results. Student’s *t*-test was used to determine levels of significance between the groups in the iTRAQ proteomics studies. All these statistical tests were performed using GraphPad Prism 7. Results are presented as mean ± SD; statistical significance is indicated as follows: \**p* ≤ 0.05; \*\**p* ≤ 0.01; \*\*\**p* ≤ 0.001.

## 3. RESULTS

### 3.1. TXR inhibits SNP stimulated nitrite in cell culture without cytotoxicity

RAW264.7 cells (10^5^ cells/ml) treated with TXR (674 μM) showed similar nitrite level with SNP stimulated controls (p<0.0001) (Fig 2A). With lower TXR levels, nitrite concentration showed an increase in trend from 25.6 μM in the highest concentration to 29.8 μM (p < 0.05) in 337 μM TXR, 30.4 μM in 168.5 μM (p<0.05). Cells treated with SNP and TXR at varied concentrations (674 - 5.26 μM) showed similar viability (Fig 2B).

**Figure 2:**
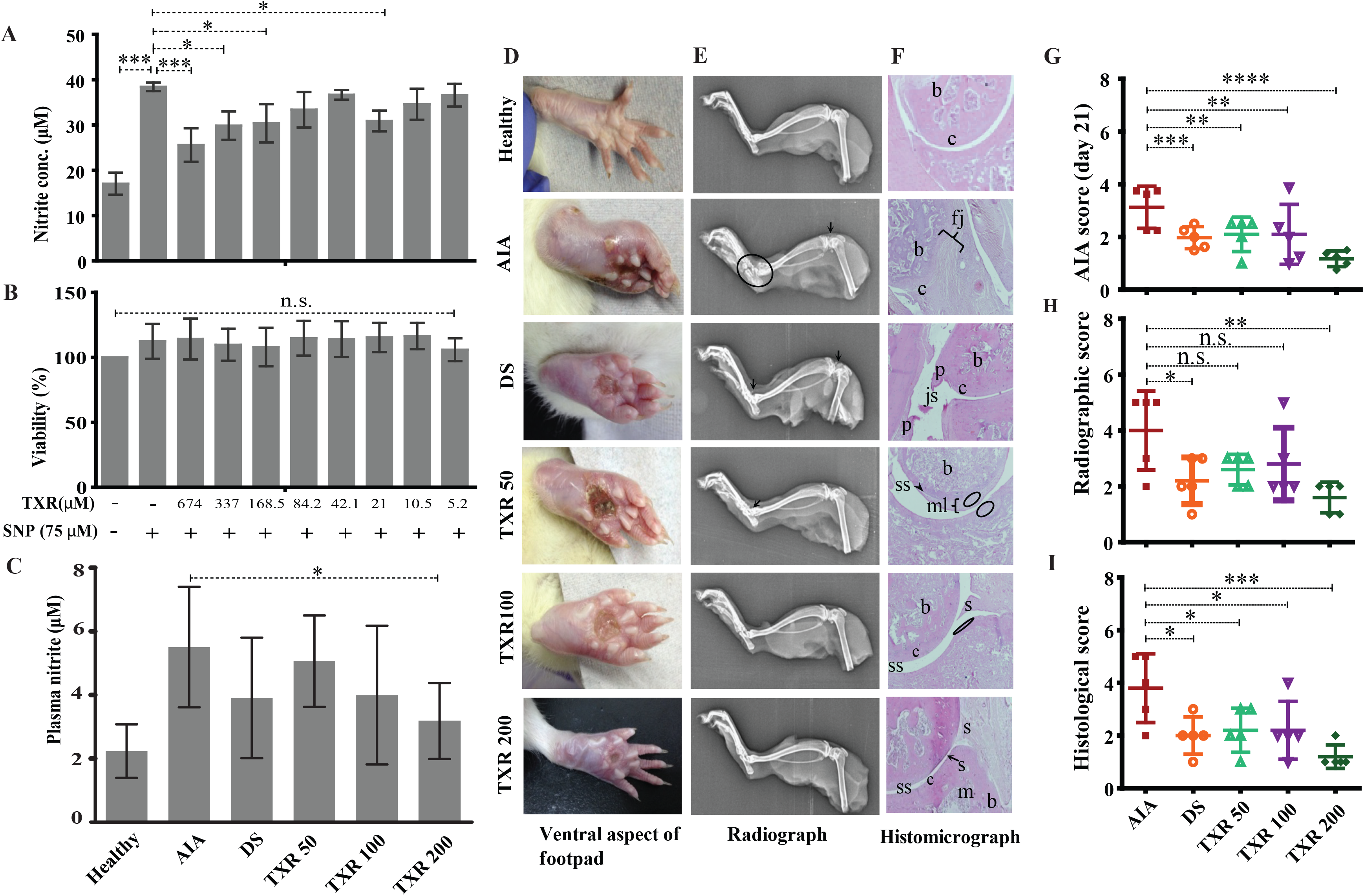
TXR has limited cytotoxicity in RAW 264.7 cell line and anti-arthritic potential in AIA rats. The released nitrite concentration in the cell supernatant (A) and viability (B) of the RAW 264.7 cells with or without TXR. C. Blood plasma nitrite content in the experimental animals using Griess nitrite assay. D. Representative pictures of the ventral aspect of foot pads. E. Radiograph of the tibiotarsal joints. Arrows indicate the osteophyte formation and oval structure mark the extent of osteolysis. F. Representative histological micrograph of the hematoxylin and eosin stained slides of rat joints. (Original magnification × 10) (js: joint space; c: cartilage; b: bone; p: pannus; s: synovium; mls: multilayered synovial membrane; fj: fused joints; m: matrix). The arrow heads indicate the damage in the synovial lining and the oval structures show neutrophils in the epiphysial cartilage. G. AIA score at day 21. H. The cumulative radiographic score. I. Histological score. AIA: adjuvant induced arthritis; DS: diclofenac sodium treated AIA; TXR 50, 100 and 200 mg/kg dose of troxerutin.

#### 3.2.1. Effect of TXR on plasma nitrite concentration

In AIA group, higher (2.5 fold) plasma nitrite level was observed with respect to healthy animals. In TXR treated groups, lower plasma nitrite levels were observed in a dose dependent manner (8.02% in TXR50, 27.49 % in TXR100 and 42.24% in TXR200) with respect to AIA group. In TXR200 group, lowest nitrite levels were observed in the plasma of AIA animals (p<0.05). TXR100 and DS showed similar effect (~29.06% decrease) on plasma nitrite levels as depicted in Fig 2C.

#### 3.2.2. Joint radiographic and histopathology analyses show protective effects of TXR

The footpad thickness at disease onset was highest and resolved during the treatment (Fig 2D). The limb roentgenograms of the experimental animals showed joint erosion with osteophyte formation; edema and soft tissue with noticeable swelling (Fig 2E). AIA group showed considerable damage with bone erosion and reduction in joint space. In TXR200 (p≤0.01) and DS (p≤0.05) groups, a significant recuperation of the joint damage were observed. AIA group exhibited drastic inflammation of the tibio-tarsal joints resulting in an increase in the thickness of bones and cartilages (Fig 2E and 2F). All the treatment groups showed reduction in osteological swelling in the tibiotarsal joints. The tibiofemoral joints were less affected as compared to tibiotarsal joints in all the groups. However, AIA and DS groups exhibited signs of damage (Fig 2E and 2F). AIA score at day 21 showed significant reduction in TXR200 group (Fig 2G). There was a marked reduction in the radiographic score of TXR50 and TXR100 but were not statistically significant (Fig 2H). In the plantar regions, reduced soft tissue swellings were observed in the TXR treated groups. The histopathololgical data of all the groups was expressed in terms of histological score (Fig 2I).

#### 3.2.3. TXR treatment suppresses disease progression in arthritis

The arthritic symptoms appeared within 18-36 hours of adjuvant immunization and the inflammatory parameters were edema, periarticular erythema, and functional decline in gait of the immunized rats (Fig 3). The gait of the arthritic animals improved substantially in all the TXR treated groups as compared to the AIA group, which was included as a parameter in arthritic score calculation throughout the treatment period (Fig 3A) and on the day of sacrifice, i.e., day 21 (Fig 2G). TXR (200 mg/kg) treatment has brought down the arthritis score close to the basal level (healthy) (Fig. 3A) and showed visible effect in the hind footpads of the rats (dorsal view, Fig 1C and ventral view, Fig 2D). Significant reduction of mean arthritic score was observed in a dose-dependent manner from day 6 (dorsal view, Fig 1C and ventral view, Fig 2D). AIA score pattern was found to be similar in TXR100 and DS groups whereas in TXR50 positive effect was observed by day 12. The arthritis ameliorating potential of TXR at 50 and 100 mg/kg was found to be comparable to DS treatment based on appearance of secondary lesions in the footpad and tail base. A marked decline in AIA score in TXR200 from day 6 (p ≥ 0.05) to day 21 (p ≤ 0.0001) was observed (Fig 3A). Dose-dependent remissions of lesions in the subplantar region were observed in TXR treated groups. TXR200 group showed rapid wound healing and erythema with respect to TXR100, TXR50 and DS groups. AIA score, on the experiment termination day (day 21), showed highly significant improvement (p<0.0001) in TXR200 followed by DS (p=0.0006) and TXR100 and TXR50 groups (both p =0.002).

**Figure 3.**
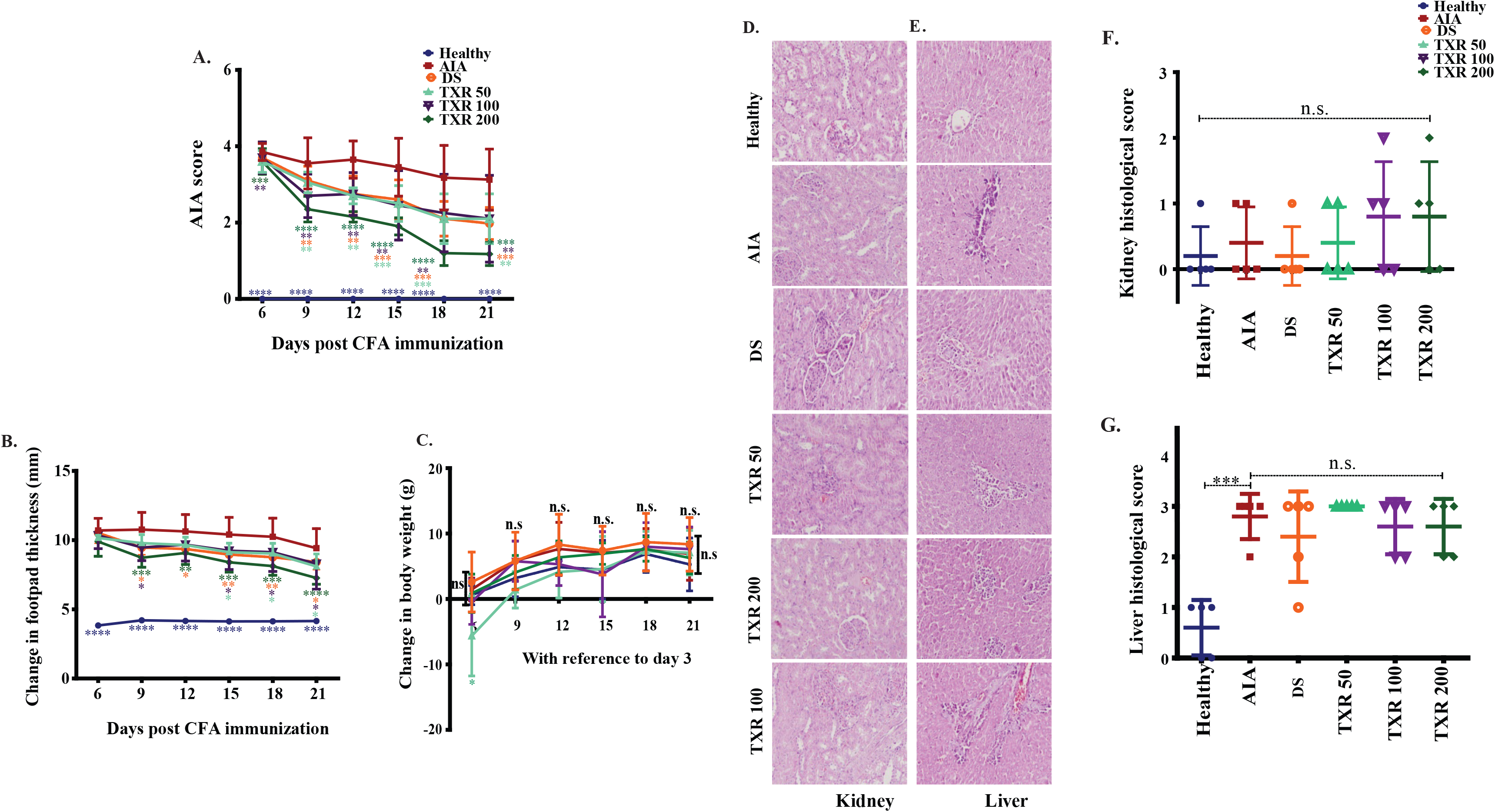
TXR treatment suppresses arthritis score in adjuvant-induced arthritis rats in a dose dependent manner. The arthritis score (mean ± SD) (A), change in foot pad thickness and (B) change in the body weight, (C) shows a trend with TXR administration. Photomicrographs of hematoxylin and eosin stained histological slides of kidney (D) and liver (E) of rats of different study groups. The histological scores of kidney (F) and liver (G) were derived from analysis of different parameters. n.s.: not significant at 95% confidence and *p*-value of less than 0.05 was considered as significant. \**p* <0.05, \*\**p* < 0.01, \*\*\**p* < 0.001, \*\*\*\**p* < 0.0001.

#### 3.2.4. Reduction in footpad thickness upon TXR treatment

Significant reduction of mean footpad thickness in TXR treated groups was observed in a dose-dependent manner (Fig 3B). In TXR200, a continuous reduction in swelling as compared to the AIA group from day 9 till day 21 was observed. TXR100 and DS exhibited similar trend throughout the experimental period while TXR50 showed least inhibitory effect.

### 3.3. TXR has minimal adverse effect in the treatment of arthritis

#### 3.3.1. Effect of TXR on body weight gain

All experimental animals showed normal weight gain during the experimental period. The AIA and healthy animal groups showed similar weight gain till day 21. Body weight gain in the TXR50 group followed a different trend than the other experimental groups (Fig 3C). TXR100 and TXR200 groups showed insignificant weight change with respect to the AIA or healthy group. This can be attributed to the beneficial effects of TXR overcoming its adverse effects due the lower dose.

#### 3.3.2. Effect of TXR on liver and kidney histology

We observed the absence of liver steatosis, sinusoidal dilatation, Kuppfer cell hyperplasia, apoptosis, and necrosis in liver tissues from TXR treated animals (Fig 3D, 3E). The presence of eosinophils in the portal tracts and sinusoids suggested drug-induced liver injury (DILI). TXR treated groups showed insignificant difference in the liver and kidney histological scores when compared to AIA group (Fig 3F and 3G). Based on the liver histological score, AIA induction contributes to liver damage (Fig 3E and 3G). With the administration of DS or TXR, the liver histological score remains similar to the AIA group. At any given dose, TXR did not improve AIA-induced liver or kidney damage (*p* >0.05) when compared to AIA.

### 3.4. Effect of AIA and TXR treatment on joint homogenate proteome

A total of 434 joint proteins were identified in the experimental groups (Fig. 4A and Table S1). Positive correlation was observed in the biological replicates of AIA (R^2^ = 0.569), TXR (R^2^ = 0.597) and DS (R^2^ = 0.512) groups (Fig S2A, S2B, and S2C). A set of 65 proteins showed deregulation (fold change ≥1.5) in AIA with respect to healthy group (Fig S2A). Forty nine (49) proteins out of 65 were found up-regulated while 16 were down-regulated in AIA group with respect to healthy controls (Table S2). When compared with AIA, in TXR treated groups, 11 (9↑ and 2↓) proteins with ≥1.5 fold change (Table S3), while 27 proteins were found to be significantly (*p* ≤ 0.05) deregulated (Table S4). When DS group was compared with AIA, 19 (7↑ and 12↓) proteins were ≥1.5 fold deregulated and 17 proteins were significantly deregulated. Three proteins (complement component 9, C-reactive protein and α-1, β-glycoprotein) were found to be significantly differentially expressed and are known inflammatory mediators playing critical roles in the arthritis pathogenesis (Fig S3A). Two important inflammatory mediator proteins (C-reactive protein and adenylate-kinase isoenzyme-1) showed deregulation in DS group (Fig S3A). A set of 5 proteins (AAT, T-kininogen, vimentin, nucleophosmin, and desmin) were sufficient to classify the AIA diseased groups from the healthy group and were selected for further validation. We have also compared the differential expression of proteins in TXR and DS treated groups with the healthy animals to find that 28 (↑24 and ↓4) proteins in TXR (vs healthy group) (Table S5), while 87 (↑76 and ↓11) proteins in DS (vs healthy group) were ≥1.5 fold changed.

**Figure 4.**
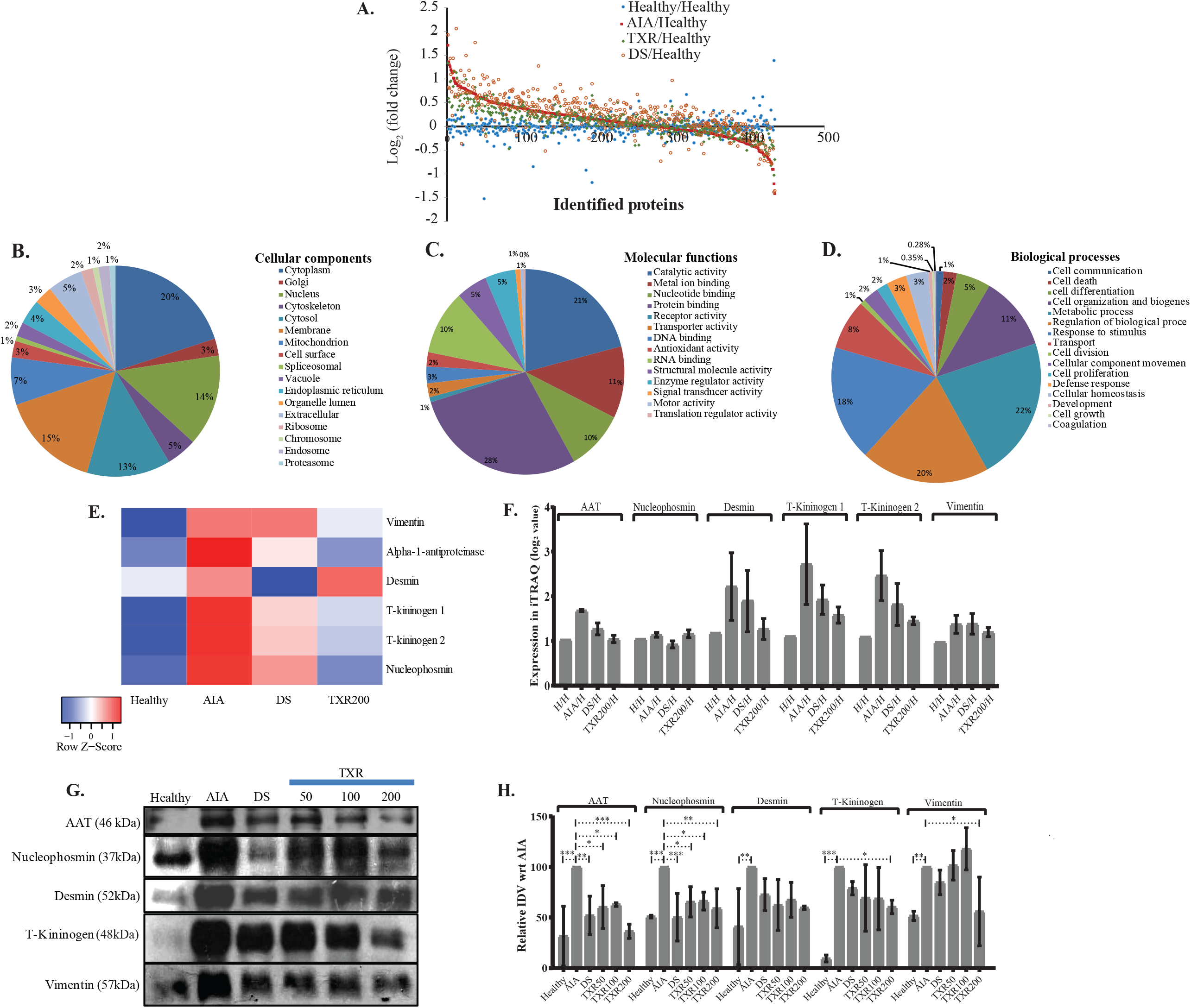
Global proteomics assay of tissue joint homegenate proteins (n=434) in experimental study groups. (A) The scatter plots showing all identified proteins and their fold change values in the experimental groups. All identified joint proteins were categorized according to cellular components (B), molecular functions (C) and biological processes (D). (E) Heatmap of important deregulated proteins that classified the study groups correctly. The color and its intensity explain the fold change values. (F) The iTRAQ ratios of the important deregulated proteins as observed in different study groups. (G) Western blot analysis of important markers molecules (alpha-1-antitrypsin, desmin, nucleophosmin, T-kininogen, and vimentin) and their relative integrated densitometric values (IDV) of Western blot bands (H). The IDV of AIA was put constant as 100% against the other experimental samples.

### 3.5. Functional categorization of the iTRAQ identified proteins

The identified proteins from iTRAQ experiment were functionally annotated according to the cellular location, molecular function and biological processes (Fig 4B, 4C and 4D). Majority of these proteins were of cytoplasmic (20%), membrane (15.4%), nucleus (14.9%), cytosolic (12.9%) and mitochondria (7.25%) origin (Fig 4B). On the other hand, the identified proteins are involved in more than one molecular function and found to possess simple protein activity (28.4%), catalytic activity (20.9%), metal ion activity (11.3%), RNA binding (10.2%), and nucleotide activity (9.5%) (Fig 4C). It was observed that in biological systems, 27.5% of these proteins are primarily involved in metabolic processes, 25.8% represent different biological regulations, 22% proteins are expressed in response to different stimuli and 14.2% are involved in cell organization and biogenesis while the rest are transport proteins (2%) (Fig. 4D).

### 3.6. Relative expression of proteins in iTRAQ and Western blot experiment

The fold change values of identified important set of proteins from the iTRAQ experiments (Fig. 4E and 4F) were monitored in independent samples using Western blot analysis (Fig 4G, 4H and S3B). For example, T-kininogen, a rodent-specific inflammatory mediator, increased (~15.4 fold) with arthritis induction and significantly reduced in TXR200 to 2.8 fold. DS, TXR100 and TXR50 groups showed similar levels of T-kininogen as observed in AIA group. Abundance of both, T-kininogen-1 and T-kininogen-2 were found to be similar in mass spectrometry and in the Western blot data. TXR200 treatment reduced both T-kininogen-1 and T-kininogen-2 levels as observed in iTRAQ analysis. Marked reduction in vimentin level was evident in TXR200 treatment whereas TXR50, TXR100 and DS did not show considerable change in vimentin as compared to AIA. Higher abundance of AAT was observed in AIA compared to healthy and with DS treatment, the expression was reduced. Level of AAT showed reduction with TXR treatment in a dose dependent way. A similar trend was observed in desmin and nucleophosmin level. With respect to AIA group, lower nucleophosmin level was observed in DS treated group.

### 3.7. The arthritic PPI network follows hierarchical scale-free topology composed of the modules at different levels of hierarchy

From the primary network of the identified proteins was constructed using the interactome network of 434 proteins, the physical interacting PPI network of 434 proteins with 434 nodes and 13,316 edges (Fig. 5A). This primary arthritic network upon analysis showed that power law distributions for the probability of node degree distribution, *P*(*k*), clustering coefficient *C*(*k*) with *negative exponents*, and neighborhood connectivity distribution *C*_*N*_(*k*) against degree (*k*) with *positive exponents* were followed (Equation 9) (Fig. 5B)^26^. This power law feature shows that the network exhibited hierarchical-scale free behavior with the system-level organization of communities. Detection of the sub-communities and communities at different levels of organization was possible with the use of Louvain modularity optimization method^21^ as shown in Fig 5C. As a result, 122 communities and smaller communities were found, and 6 of them could reach up to the bottom *motif* level.

**Figure 5.**
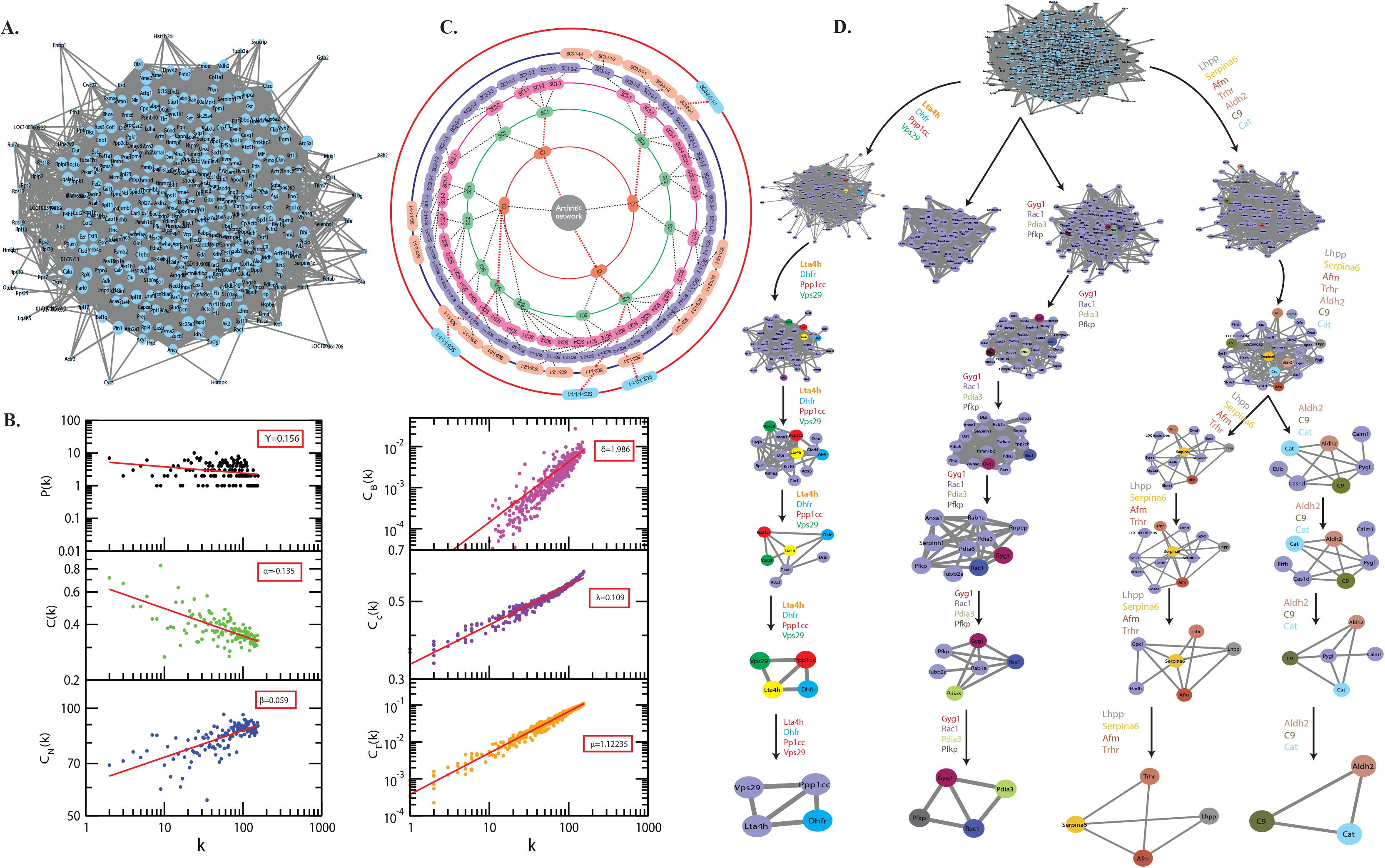
A system level organization of protein network in arthritis model. (A) The figure shows all the networks comprising of all 434 proteins; this is the first level of protein network. (B) The plots of LCP-correlation is a function of CN for each modules/ submodules (plots correspond to each module/sub-module of the network) of C9 path. This also contains the plots of PH and PLCP as a function of the level of organization. (C) Organization of sub-modules and modules at different levels as indicated by concentric circles while the arrows indicate sub-modules built from previous modules leading to the identification of key regulators of arthritis network.(D) The modular path of the key regulator proteins from complete network to motif with the structures of modules/ submodules at different stages of community finding. This leads to finding out three sub-modules through which the first four leading hubs passed through. These leading hubs comprise of the fifteen *in silico* key regulators and the probability distribution of the latter is a function of the degree of organization.

Communities at first hierarchical level exhibited the power law distribution for *P*(*k*) and C(k) against degree distribution with negative exponents demonstrating further system-level organization of the modules (Equation 9). C_N_ (k) exhibits the power law against degree *k* with a positive exponent (*β* ~ 0.05, 0.13 and 0.14, respectively) (Fig 5D). This specifies the assortivity nature of the modules reflecting the possibility of the formation of rich-club and the hubs play a very important role in the maintenance of network stability and properties^26^.

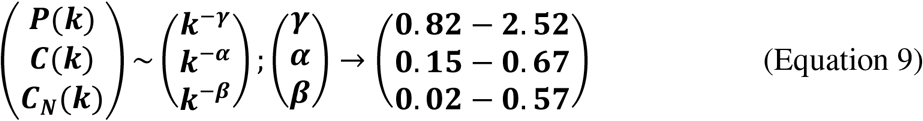

### 3.8. The fifteen(15) novel key regulators of the network are considered to be its backbone

There is a very important role of the nodes in the information processing within a network and this is well assessed by the centrality measures such as the C_B_, C_C_, and C_E_. These are the different topological properties that determine the signal transmission efficiency of a network ^31^. In the network and modules of arthritis at first hierarchical level, these factors also demonstrated power law as the function of degree (k) with positive *exponents*, with increase in degree of nodes the centralities tend to get increased (Equation 10) (Fig 5B). The values of the exponents of C_B_, C_C_, and C_E_ for the first level of network organization are found to be δ=1.986, λ=0.109 and μ=1.12235.

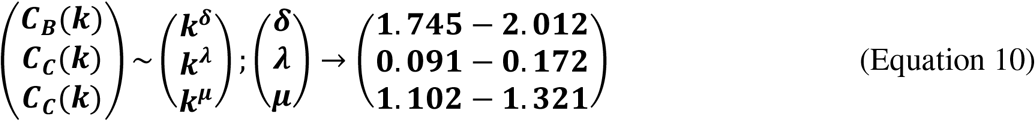

Thus, there is an increase in the signal processing efficiency with higher degree nodes emphasizing the significant roles of these nodes in the flow of information, regulation and stabilization of the network. Therefore, the *hub* proteins must have played a significantly large influence in the network regulation and pathogenesis of arthritis. The 122 modules of proteins were taken into consideration to find out the most important proteins as they were present at each topological level resulting in the identification of the most high-ranking key regulator proteins in the arthritic network. After tracing *hubs* at every topological level, 15 proteins (*C9, Aldh2, Pdia3, Serpina6, Afm, Gyg1, Ppp1cc, Pfkp, Dhfr, Cat, Trhr, Vps29, Lta4h, Rac1, Lhpp*) (Table S6) were established as the backbone of the entire network. The key regulators which form the *motifs* with their partners (Fig 5D) might be instrumental in the network integrity, optimization of signal processing, dynamics, maintaining the stability and most importantly regulation of the network. Our community finding method confirmed that all the 15 key regulators are interacting with each other (Fig 6A). This PPI was later validated using the database of STRING 10.0 (Fig 6B).

**Figure 6.**
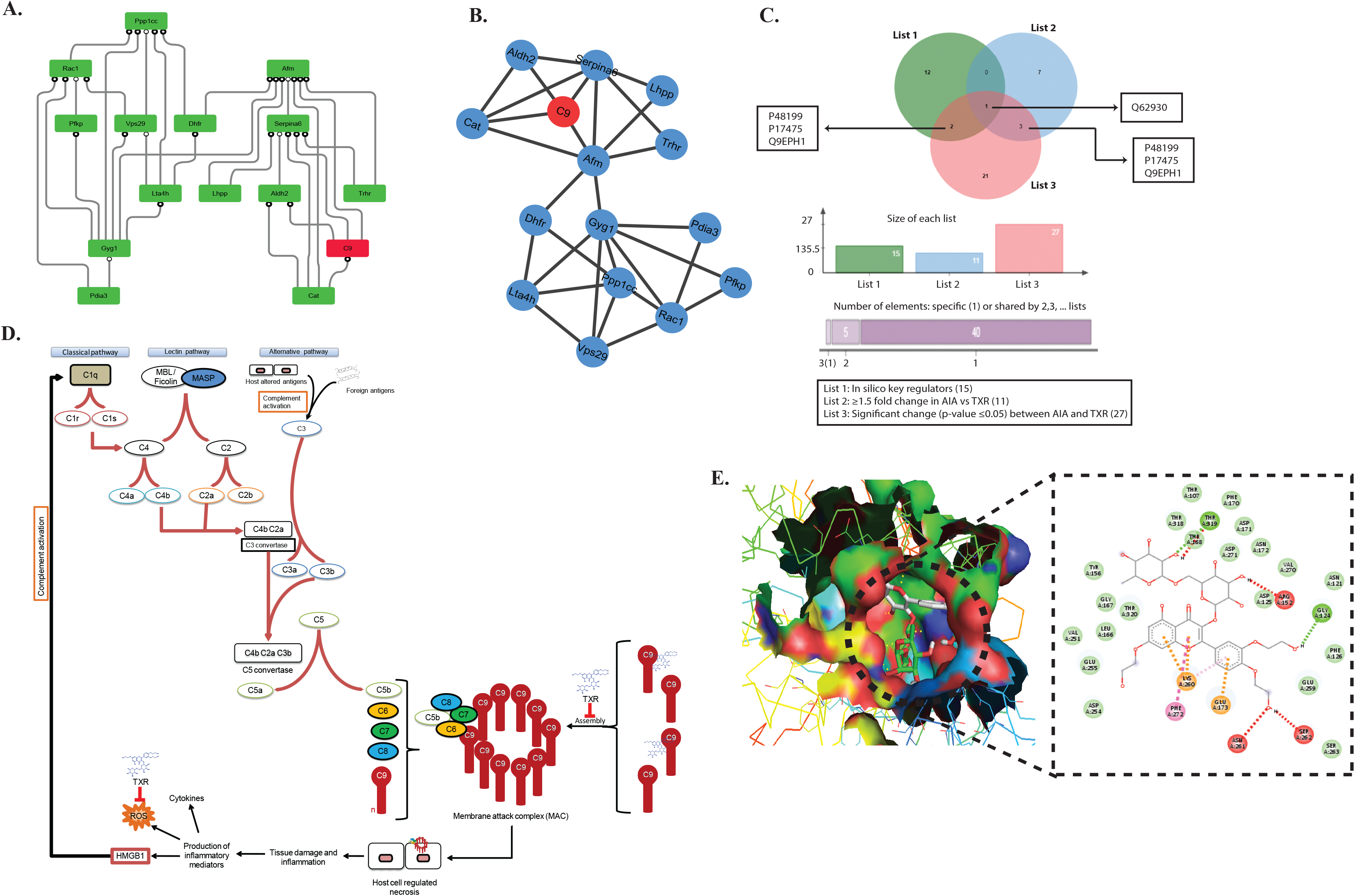
Mechanism of action study of TXR. (A) Protein-protein interaction (PPI) of all the fifteen key regulator proteins computationally derived through the community finding method. (B) String interaction network of the fifteen *in silico* key-regulator proteins. The PPI networks were constructed using STRING 10.0 with a medium confidence level (0.4) and all available prediction methods. (C) Venn diagram showing C9 as the common protein in all the lists of proteins, viz. list 1 = 15 proteins (*in silico* key regulators), list 2 = 11 proteins (≥1.5 fold expressed proteins in AIA when compared with the TXR group), and list 3 = 27 proteins (significantly differentially expressed (*p* ≤ 0.05) proteins between AIA and TXR). (D) Probable step of TXR action on C9 based membrane attach complex formation and its role in inflammation and arthritis. (E) Molecular docking of C9 protein with TXR. Expanded region of the docking site shows the interacting amino acids of C9 protein with the TXR molecule in the Ligplot.

### 3.9. C9 as the most common protein in different groups

We found four proteins (C9, protein disulphide isomerase A3, thyrotropin releasing hormone receptor, and isoform gamma-2 of serine / threonine – protein phosphatase 1) as common from list of *in silico* key regulators (n=15) and the list of ≥1.5 fold changed proteins in AIA vs healthy (n=65) (Fig S4A). Whereas, 2 proteins (adenylate kinase isoenzyme −1 and C-reactive protein) were found to be common when the list of ≥1.5 fold changed proteins in AIA vs DS (n=19) and the list of significantly (*p* ≤ 0.05) deregulated proteins in AIA vs DS (n = 17) were matched (Fig S4B). The Venn diagram presents the common list of proteins from *in silico* key regulator proteins (n=15), ≥1.5 fold changed proteins in AIA vs healthy (n=65), ≥1.5 fold changed proteins in AIA vs TXR (n=11) and significantly changed proteins between AIA vs TXR (n=27) (Fig S4C). C9 was identified as the most common protein from these 4 lists. This signifies the importance of C9 as the molecular target of TXR. We did not find and common proteins from the five lists (except n=65 list) (Fig S4D).

### 3.10. Molecular interaction of key regulators with TXR through *in silico* docking

The combination of these distinct methods viz. iTRAQ differential proteomics and *in silico* network analysis studies helped to narrow down our search to the very specific protein with probability of being the target of TXR, i.e., the complement component 9 (C9) (Fig 6C). The role of C9 is critical for the formation of membrane attack complex (MAC) resulting in tissue injury which further activates the entire complement pathway (Fig 6D). All these compounds were found to position it in the deep cavity of the proteins which show several close interactions to their catalytic residues (Fig S5) along with C9 (Fig 6E). Here, several residues of binding pocket are forming strong hydrogen bonds with the compound TXR in addition to several Van der Waals and other weak interactions to properly hold it in the binding cavity of the proteins. So, we propose TXR as a potential scaffold which can be used in the development of potential inhibitors of this protein.

Molecular docking studies with C9 showing highest binding affinity (−8.9 kcal/mol) of TXR with interacting residues of ALA86, ALA201, GLY193, GLU77, PRO85 (Table S6, Fig 6E) followed by the other 14 key regulators Aldh2 (−7.3 kcal/mol), Pdia3 (−8.0 kcal/mol), Serpina6 (−5.9 kcal/mol), Afm (−7.9 kcal/mol), Gyg1 (−6.7 kcal/mol), Ppp1cc (−6.7 kcal/mol), Pfkp (−7.3 kcal/mol), Dhfr (−7.5 kcal/mol), Cat (−7.1 kcal/mol), Trhr (−6.6 kcal/mol), Vps29 (−7.2 kcal/mol), Lta4h (−8.5 kcal/mol), Rac1 (−6.3 kcal/mol), Lhpp (−6.4 kcal/mol) (Table S6 and Fig S5). Proteins associated with inflammatory processes such as acute phase plasma proteins or others may indicate an intervention by TXR in the disease progression. Therefore, to investigate the effect of therapeutic compound on the expression of plasma proteins in the arthritic rats was of importance in our earlier study also^37^. We used the network theoretical approach which has considered the *hubs*, *motifs* and *modules* of the network with equal emphasis for the identification of key regulators or the significant regulatory pathways preventing any bias towards the overrepresented *hubs* or *motifs*. A relationship between *hubs, motifs* and *modules* was established and the network used all proteins associated with the disease instead of merely the manually curated datasets’ usage. In conclusion, the *hubs* with highest degree were identified, fifteen (15) were considered as the novel key regulators.

## 4. Discussion

The autoimmune disease, RA is considered to be an incurable and difficult to manage disease, identification of pharmacological compounds with minimum side effects is a way forward to for its management. The current pharmaceutical solutions involve disease modifying arthritis drugs which work in ~60% of the cases and in certain populations can lead to adverse reactions like pneumonia, tuberculosis, and interstitial pneumonitis. Identification of predictive markers might facilitate to develop personalized therapy to gain optimum treatment benefit in RA patients. In this study, we monitored the anti-arthritic potential of a phytochemical i.e., TXR and investigated its probable mechanism of action by identifying its target proteins in the joint homogenate.

In our *in vitro* studies, TXR has significantly reduced the nitrite levels with negligible effects on cell viability up to a concentration of 674 μM as shown by MTT assay. It is widely reported that TXR is a phytochemical with proven health benefits. It also has a very low toxicity (LD_50_ of 27160 mg/kg body weight in rat) and safe for human use^38–41^.

We used commonly adopted AIA animal model and observed manifestation of the RA disease parameters. With the TXR treatment, there was a significant reduction of radiographic and histological scores in all the AIA study groups receiving treatment in a dose-dependent manner. Significant decrease in footpad thickness was observed and similar body weight gain was noticed in the TXR treated AIA groups. Based on the histological analyses of liver and kidneys, it seems that the damage caused by AIA was not corrected with TXR or DS treatment. We used DS as a control drug and other antiarthritic drugs could be used for comparative analysis in subsequent studies. A set of 65 proteins was found to be dysregulated in the joint homogenate of AIA group as compared to the healthy group of which 11 proteins showed positive correlation with TXR treatment. Many of these deregulated proteins participate in inflammation, arthritis, autoimmune disorders and cancer. Upon TXR administration, many dysregulated proteins at the time of disease onset showed time dependent reversal to the basal level. A set of 5 proteins (AAT, nucleophosmin, desmin, T-kininogen and vimentin) were sufficient to classify the AIA diseased groups from the healthy group and were thus selected for Western blot validation. Abundance of these proteins were probed in the joint tissues of independent sample sets using Western blotting experiment and corroborated the mass spectrometry findings.

The PPI study of the identified protein sets and the protein community finding by *in silico* network analysis identified a set of key regulator molecules^42–45^. The constructed primary PPI arthritis network tracked from the level of *hubs* up to the *motifs* resulted in the detection of key regulators (*hubs*) from a total of 434 identified proteins. Employing Newman and Girvan’s method ^28^ for community finding with equal importance to the *modules*, *motifs* and *hubs* of the network, resulting in the identification of 15 novel key regulators (*C9, Aldh2, Pdia3, Serpina6, Afm, Gyg1, Ppp1cc, Pfkp, Dhfr, Cat, Trhr, Vps29, Lta4h, Rac1, Lhpp*) which are known markers involved in some other diseases also. Besides, these key regulators have other interacting partners at the level of *motifs* which may also have importance in arthritis pathogenic mechanism thus establishing them also as candidate disease regulators. Combining the mass spectrometry and *in silico* analyses, the complement component 9, C-reactive protein, catalase, aldehyde dehydrogenase, α-1-antiproteinase and α-1B glycoprotein (AAG) were selected as the common key regulators.

Upon TXR treatment, the C9, AAG and catalase abundance were decreased almost to the basal levels without significant alteration in CRP, a non-specific indicator of inflammation. High aldehyde dehydrogenase (ALDH) activity has been reported in osteoarthritis patients’ chondrocytes and elevation of Serpina1 were also observed in the joint homogenates of the AIA group^46^. ALDH and AAT are known to have tissue protective properties in arthritis. AAT is an acute phase protein, possessing immunoregulatory and anti-inflammatory functions independent of antiproteinase activity. In the collagen induced arthritis (CIA) model, higher AAG level has been reported^47,48^. C9 is one of the multimeric components of the terminal stage of all the complement pathways, resulting in the MAC which initiates cellular lysis at the target tissue i.e. chondrocytes of the cartilage in the synovial joints. C9 may be responsible for the cartilage degradation leading to joint tissue damage in the autoimmune and reactive arthritis conditions. We have found that TXR disrupts the MAC mediated complement pathway (Fig 6D), however, further investigation at the biophysical level will be beneficial.

Activation of the entire complement cascade, results in the formation of MAC which initiates pro-inflammatory responses causing autoimmune diseases^49^. So, TXR ameliorates arthritis through inhibition of the complement-mediated autoimmune pathogenesis. There have been many successful attempts for the containment of the complement mediated pathogenesis in arthritis by targeting various components of all the three major complement pathways^50^. Lappegård and coworkers describe various inhibitors of the complement component pathway, however, to the best of our knowledge there has been no inhibitor or blocker reported for the C9 component of MAC. Through our present study, we propose that TXR is a potent inhibitor of the MAC which is a complex of transmembrane proteins composed of C9 subunits. MAC is responsible for channel formation across the plasma membrane of the target cell resulting in its lysis. The channel thus formed allows the inflow of several ions (Ca^2+^, Na^+^) causing endosmosis followed by necrotic cytolysis, thus releasing the cytokines and other related inflammatory mediators into the milieu. So it may promote the production of interleukin 1 beta (IL-1β) through Nod-like receptor protein 3 (NLRP3) inflammasome thus acting as an immune stimulating factor^51^. MAC mediated necrosis of the target cells (chondrocytes, osteocytes and synoviocytes) release inflammatory mediators like cytokines, high-mobility group box 1 (HMGB1) resulting further joint damage^52^. The cytokines, in turn, stimulate the other cells to activate the complement pathway^53^, thus aggravating the inflammatory response manifold. Extracellular HMGB1 binds to C1q thus activating the classical complement pathway followed by the reformation of MAC^54^. This pathway is probably one of the leading mechanisms in joint dysfunction in autoimmune arthritis. If unchecked, this vicious cycle goes on to further debilitate the synovial joint tissues. It also plays an important role in tissue degeneration, neuroinflammation as well as arthritis^55,56^.

Molecular docking study showed that TXR has high affinity towards C9, so it seems that TXR will be even more effective to cause steric hindrance resulting in the disassembly of the multimeric C9 in the MAC. This may inhibit the inadvertent cellular lysis and joint damage in case of sterile inflammation or related pathogenesis^57^. So, TXR may inhibit necrosis mediated cell death by blocking C9 involvement in MAC formation. Since, TXR is well tolerated; inhibition of its possible molecular targets may have minimal or no adverse effects. TXR has been widely reported to possess hepatoprotective^58,59^ as well as renoprotective^60^ properties which can now be attributed to the MAC inhibition thus protecting the hepatic and renal cells from xenobiotic mediated lysis. TXR is also a known antioxidant and inhibits oxidative stress mediated cellular apoptosis^61^ and may protect the synovial joints alleviating severe damage in AIA and possibly in rheumatoid arthritis patients.

Natural products that could inhibit production of chemokines and cytokines and modulate osteo-immune cross-talk could be useful in the treatment modalities for RA. These molecules may vary through many other inflammatory mediators such as NF-κB, MAPK, and STAT3, etc. It can also be inferred that the effect of TXR might have resulted in the inhibition of HMGB1 thus reducing inflammation^62^. The effect of TXR on the immune systems and their effector molecules needs additional studies.

The identified candidate biomarkers in responses to the drugs for RA suggest new modalities of anti-arthritic treatment. However, the observations from experimental animal model need further validation of the identified target to understand the perturbed molecular details involved in this disease pathophysiology. Understanding the molecular pathway will be useful to identify alternate druggable targets for new molecule discovery. Our results demonstrated that TXR has therapeutically beneficial effects on the experimental arthritis animal model and may be useful as a potential treatment for RA in humans after appropriate clinical trials.

In conclusion, our quantitative proteomics approach demonstrated the anti-arthritic properties of a phytochemical TXR and its probable interaction with synovial proteins. Combination of quantitative proteomics study supported by the robust protein community finding method provided a comprehensive tool to map probable targets of TXR and its mechanism of action. The experimental approach adopted in this study will be useful at various phases of drug discovery and validation in translational studies for various disease conditions.

## Acknowledgements

Debasis Sahu was supported by DST-SERB Young Scientist Award and the study was also funded with a start-up research grant by SERB, Government of India. Authors acknowledge Dr. Perumal Nagarajan and the staff of animal research facility in National Institute of Immunology (NII), for providing support for conducting experiments. RKN acknowledges core support from ICGEB New Delhi.

## Figure legends

**Figure S1.** (A) Scheme of iTRAQ labeling: Flowchart showing the eight-plex iTRAQ labeling and the quantitative proteomics method used for analyzing joint homogenate proteins. The iTRAQ tags 113 and 115 were used for the technical replicates of a healthy animal, while biological replicates of AIA were tagged with 114 and 116, followed by those of TXR with 117, 119 and lastly 118, 121 were used for those of DS. (B) One dimensional silver stained gel shows the protein band patterns in all study group samples.

**Figure S2.** Scatter plots showing the correlation between the protein expressions of biological replicates in the experimental groups: Scattering plots of Pearson correlation comparing two biological replicates of iTRAQ reporter ion intensities, (A) AIA1 (114/113) vs AIA2 (116/113); (B) DS1 (118/113) vs DS2 (121/113); and (C) TXR1 (117/113) vs TXR2 (119/113). The Pearson correlation coefficient (R^2^) is given for each plot.

**Figure S3.** (A) Heat map of the proteins showing both significant difference and ≥1.5 fold change between the AIA and treatment groups (both DS and TXR200). (B) Western blotting of the joint homogenate proteins of a different set of animals from each experimental group as a replicate result.

**Figure S4.** (A) Venn diagram showing 4 common proteins between 2 lists of proteins, viz. list 1 = 15 proteins (*in silico* key regulators), list 2 = 65 proteins (≥1.5 fold expressed proteins between AIA and Healthy). (B) Venn diagram showing 2 common proteins between 2 lists of proteins, viz. list 1 = 19 proteins (≥1.5 fold expressed proteins between AIA and DS) and list 2 = 17 proteins (significantly differentially expressed (p ≤ 0.05) proteins between AIA and DS).

(C) Venn diagram showing C9 as common among 4 lists of proteins, viz. list 1 = 15 proteins (*in silico* key regulators), list 2 = 65 proteins (≥1.5 fold expressed proteins between AIA and Healthy), list 3 = 11 proteins (≥1.5 fold expressed proteins between AIA and TXR groups), and list 4 = 27 proteins (significantly differentially expressed (p ≤ 0.05) proteins between AIA and TXR). (D) Venn diagram showing many sets of common proteins among 5 lists of proteins, viz. list 1 = 15 proteins (*in silico* key regulators), list 2 = 11 proteins (≥1.5 fold expressed proteins between AIA and TXR groups) list 3 = 27 proteins (significantly differentially expressed (p ≤ 0.05) proteins between AIA and TXR), list 4 = 19 proteins (≥1.5 fold expressed proteins between AIA and DS) and list 5 = 17 proteins (significantly differentially expressed (p ≤ 0.05) proteins between AIA and DS).

**Figure S5.** Molecular docking interaction studies (molecular interaction is depicted on left while Ligplot showing molecules involved in interactions on right side of each column) of TXR with the proteins identified as *in silico* key regulators. (A). TXR with complement component 9 (C9); (B). TXR with catalase (Cat); (C). TXR with aldehyde dehydrogenase 2 (Aldh2); (D). TXR with Phosphofructokinase, platelet (Pfkp); (E). TXR with afamin (Afm); (F). TXR with leukotriene A4 hydrolase (LAH); (G). glycogenin 1 (Gyg1); (H). TXR with VPS29 retromer complex component (Vps29); (I). TXR with phospholysine phosphohistidine inorganic pyrophosphate phosphatase (Lhpp); (J). TXR with protein phosphatase 1 catalytic subunit gamma (Ppp1cc); (K). TXR with Serpin family A member 6 (Serpina 6); (L). TXR with Ras-related C3 botulinum toxin substrate 1 (Rac 1); (M). TXR with Thyrotropin releasing hormone receptor (Trhr); (N). TXR with Protein disulfide isomerase family A, member 3 (Pdia3); (O). TXR with dihydrofolate reductase (Dhfr).

