## Supplementary material for "Troxerutin acts on complement mediated inflammation to ameliorate arthritic symptoms in rats": Fig S1

**Figure S1**

A.

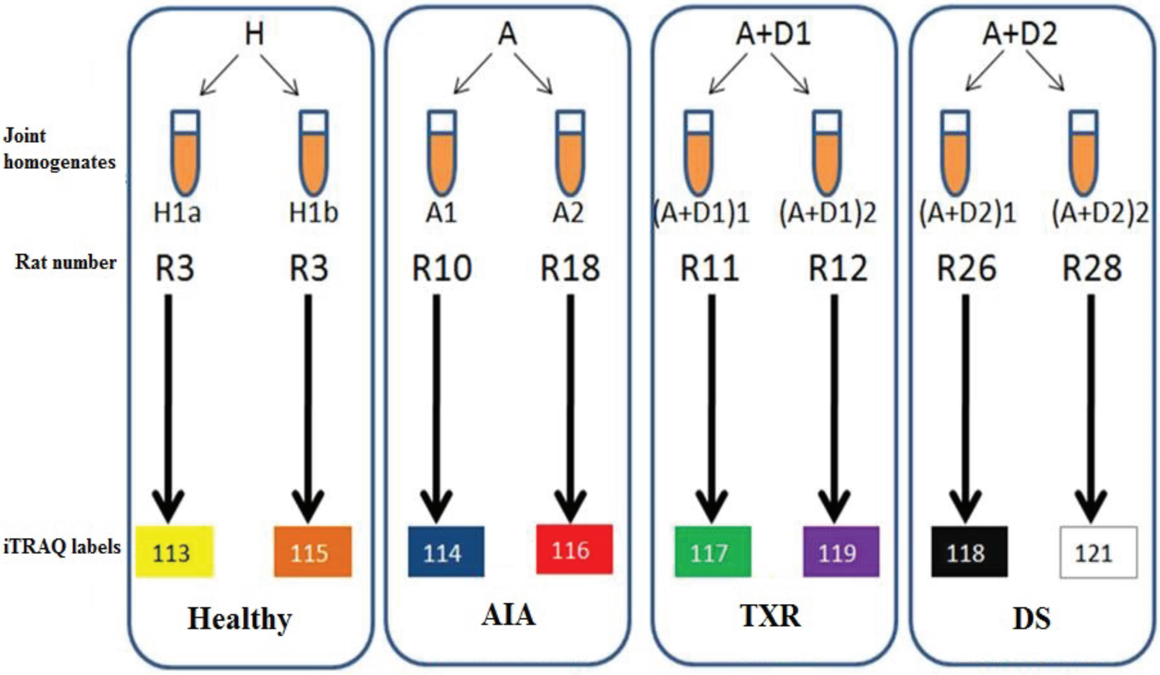


B.

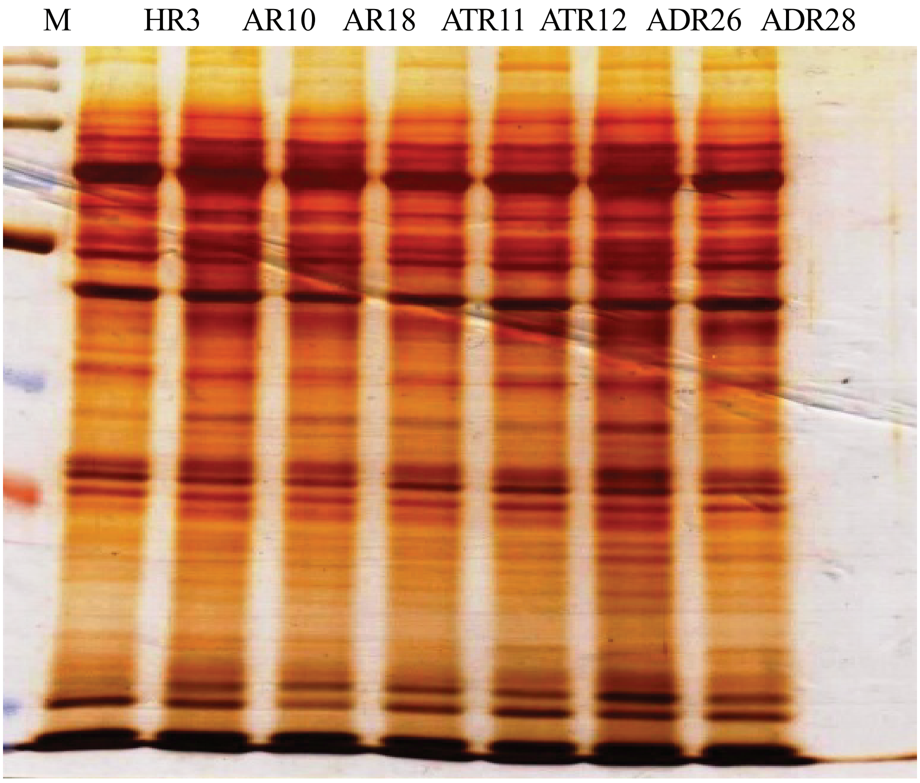


M - Marker
HR3 - Healthy
AR10 - Arthritis (no treatment)
AR18 - Arthritis (no treatment)
ATR11 - Arthritis + TXR (200mg/kg)
ATR12 - Arthritis + TXR (200mg/kg)
ADR26 - Arthritis + Diclofenac sodium
ADR28 - Arthritis + Diclofenac sodium
