## Supplementary figures and images for "Troxerutin acts on complement mediated inflammation to ameliorate arthritic symptoms in rats"

### Fig S2

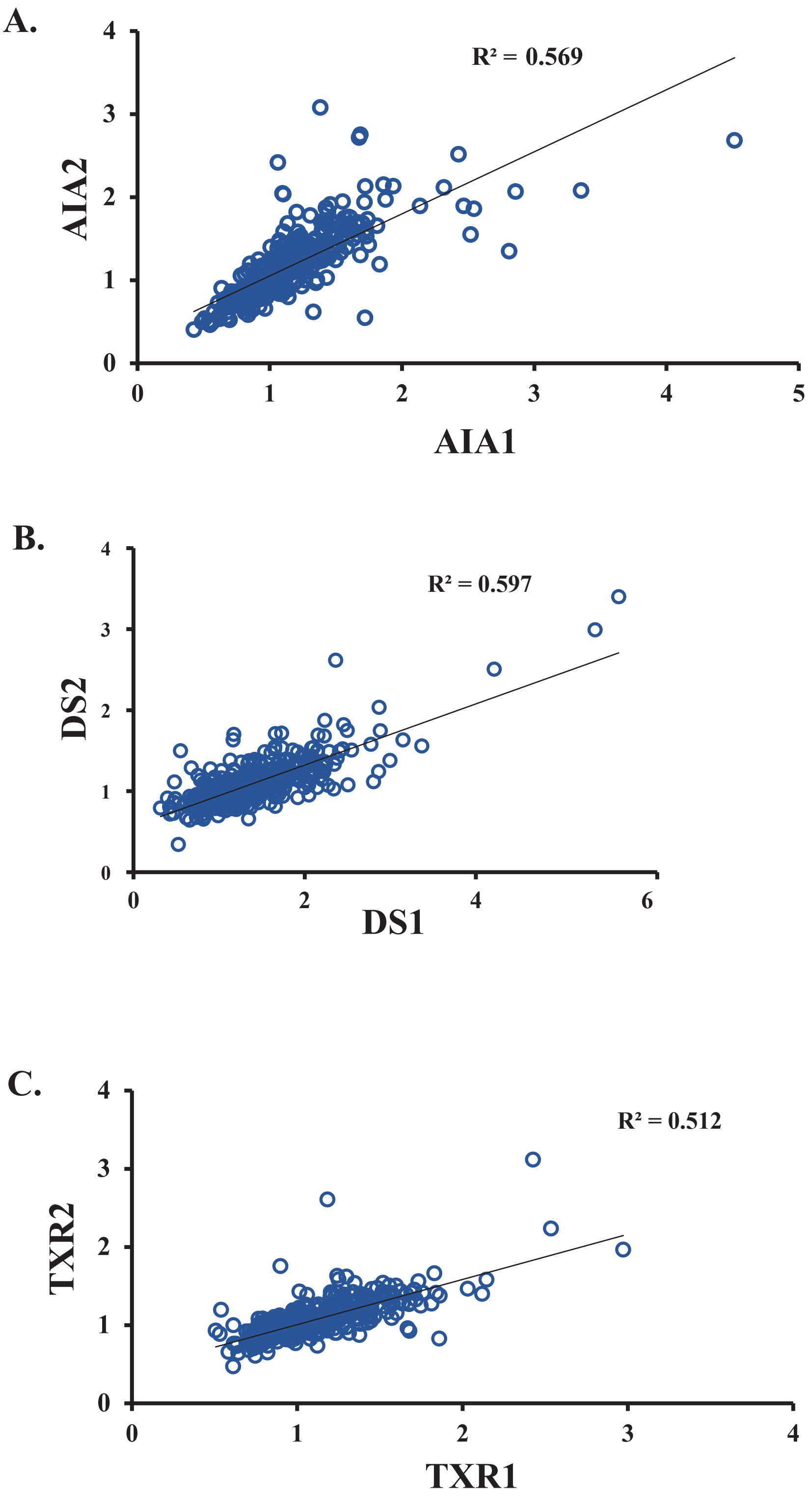

### Fig S3

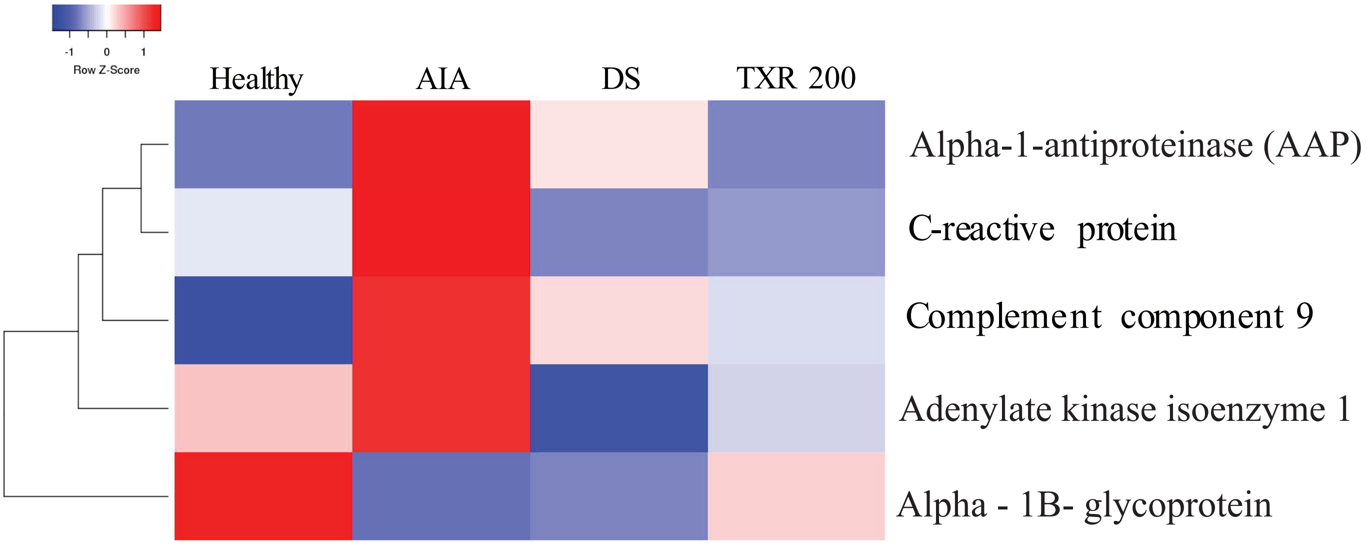

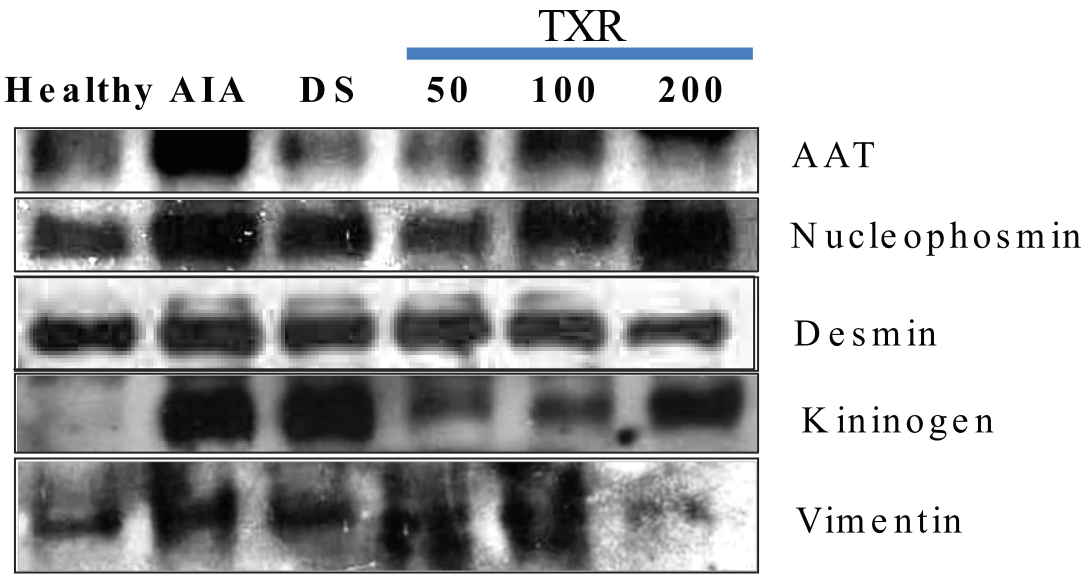

### Fig S5

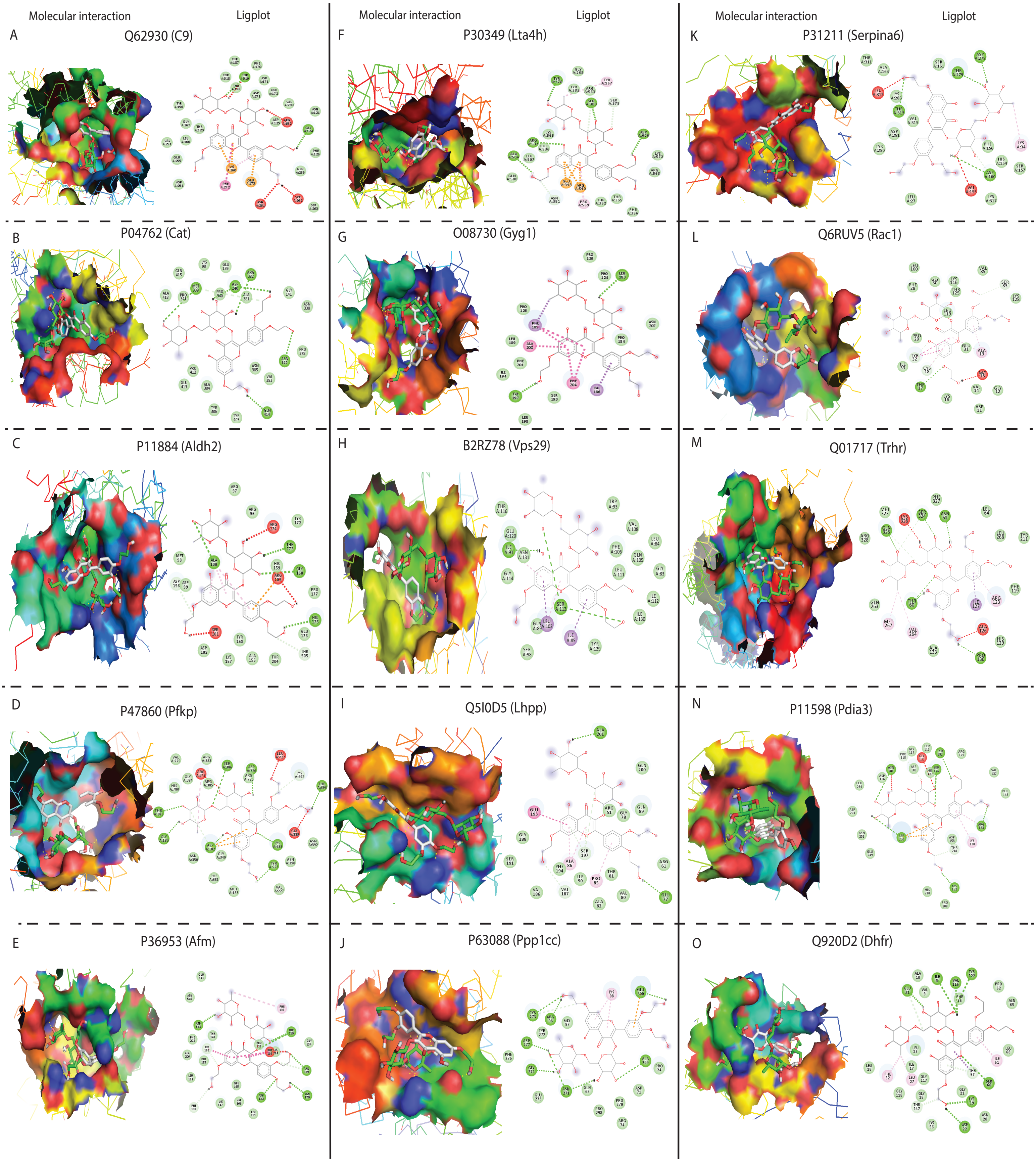
