## Supplementary material for "Troxerutin acts on complement mediated inflammation to ameliorate arthritic symptoms in rats": Fig S4

### Figure S4

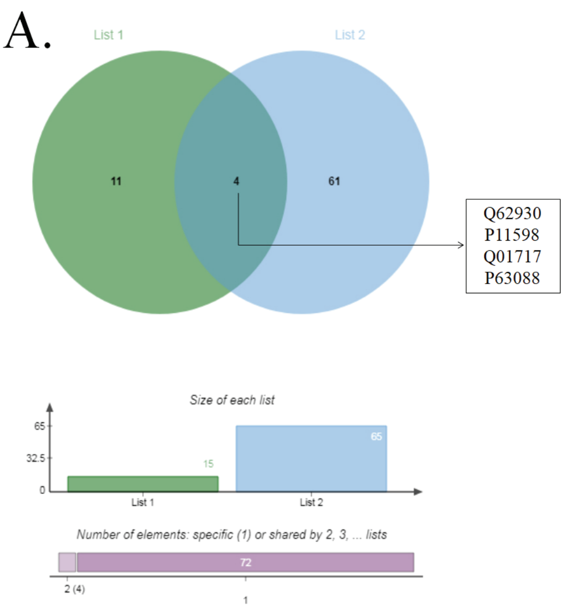


**List 1:** *In silico* key regulators (15)
**List 2:** ≥1.5 fold change in AIA vs Healthy (65)

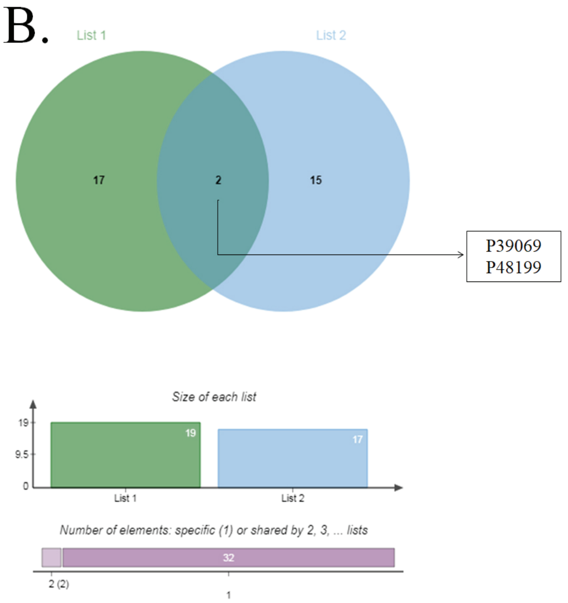


**List 1:** ≥1.5 fold change in AIA vs DS (19)
**List 2:** Significant change ($p$-value ≤0.05) between AIA and DS (17)

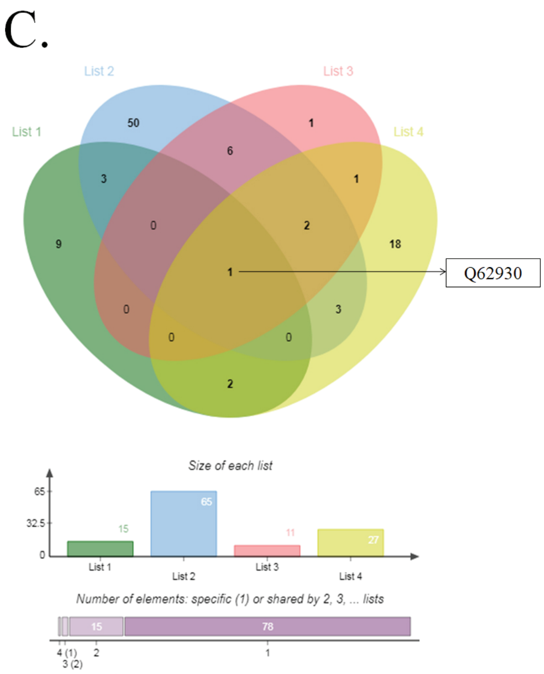


**List 1:** *In silico* key regulators (15)
**List 2:** ≥1.5 fold change in AIA vs Healthy (65)
**List 3:** ≥1.5 fold change in AIA vs TXR (11)
**List 4:** Significant change ($p$-value ≤0.05) between AIA and TXR (27)

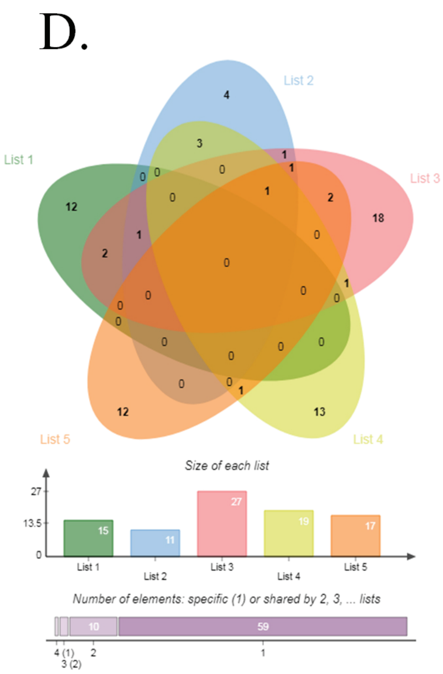


**List 1:** *In silico* key regulators (15)
**List 2:** ≥1.5 fold change in AIA vs TXR (11)
**List 3:** Significant change ($p$-value ≤0.05) between AIA and TXR (27)
**List 4:** ≥1.5 fold change in AIA vs DS (19)
**List 5:** Significant change ($p$-value ≤0.05) between AIA and DS (17)
