## Supplementary material for "Troxerutin acts on complement mediated inflammation to ameliorate arthritic symptoms in rats": Table S2, Table S3, Table S4, Table S5, Table S6

**Table S2: Identified proteins showing ≥1.5 fold change between AIA and Healthy study groups.**

| **Accession** | **Protein Name** | **Ratio (AIA/Healthy)** |
| --- | --- | --- |
| P02764 | Alpha-1-acid glycoprotein | 3.27 |
| Q62930 | complement component C9 | 2.56 |
| P01048 | T-kininogen 1 | 2.45 |
| P06302 | Prothymosin alpha | 2.38 |
| P20059 | Hemopexin | 2.29 |
| P08932 | T-kininogen 2 | 2.24 |
| Q5I0G4 | Glycine--tRNA ligase | 2.18 |
| Q62829 | Serine/threonine-protein kinase PAK 3 | 2.02 |
| P02650 | Apolipoprotein E | 2.01 |
| P20759 | Ig gamma-1 chain C region | 1.93 |
| P13084 | Nucleophosmin | 1.87 |
| P10960 | Prosaposin | 1.87 |
| P63088-2 | Isoform Gamma-2 of Serine/threonine-protein phosphatase PP1-gamma catalytic subunit | 1.86 |
| P13635 | Ceruloplasmin | 1.81 |
| D3ZHA0 | Filamin-C | 1.8 |
| P29457 | Serpin H1 | 1.8 |
| P06866-2 | Isoform 2 of Haptoglobin | 1.8 |
| Q6P686 | osteoclast-stimulating factor 1 | 1.78 |
| P50116 | Protein S100-A9 | 1.75 |
| Q63041 | Alpha-1-macroglobulin | 1.75 |
| Q6QGW5 | Steroid receptor RNA activator 1 | 1.75 |
| P02401 | 60S acidic ribosomal protein P2 | 1.74 |
| P11598 | Protein disulfide-isomerase A3 | 1.74 |
| P34058 | Heat shock protein HSP 90-beta | 1.72 |
| P24268 | Cathepsin D | 1.7 |
| P61983 | 14-3-3 protein gamma | 1.69 |
| P05942 | Protein S100-A4 | 1.68 |
| Q63081 | Protein disulfide-isomerase A6 | 1.67 |
| P80067 | Dipeptidyl peptidase 1 | 1.67 |
| P05371 | Clusterin | 1.66 |
| B2RYG6 | Ubiquitin thioesterase otub1 | 1.65 |
| P20761 | Ig gamma-2b chain C region | 1.64 |
| P17475 | Alpha-1-antiproteinase | 1.63 |
| Q66HD0-1 | Endoplasmin | 1.62 |
| Q5M7U6 | Actin-related protein 2 | 1.62 |
| Q68FR6 | elongation factor 1-gamma | 1.61 |
| P07895 | Superoxide dismutase [Mn], mitochondrial | 1.6 |
| P62853 | 40S ribosomal protein S25 | 1.59 |
| Q6P9V9 | Tubulin alpha-1B chain | 1.56 |
| P62919 | 60S ribosomal protein L8 | 1.54 |
| P00786 | Pro-cathepsin H | 1.53 |
| P20767 | Ig lambda-2 chain C region | 1.53 |
| Q9JJ54 | heterogeneous nuclear ribonucleoprotein D0 | 1.53 |
| P18420 | Proteasome subunit alpha type-1 | 1.52 |
| Q63416 | Inter-alpha-trypsin inhibitor heavy chain H3 | 1.52 |
| P20760 | Ig gamma-2A chain C region | 1.51 |
| Q6B345 | protein S100-A11 | 1.51 |
| Q6PDV7 | 60S ribosomal protein L10 | 1.5 |
| Q6IFV3 | Keratin, type I cytoskeletal 15 | 1.5 |
| P09605 | Creatine kinase S-type, mitochondrial | 0.66 |
| P11980-1 | Pyruvate kinase PKM | 0.66 |
| P02625 | Parvalbumin alpha | 0.65 |
| P06214 | Delta-aminolevulinic acid dehydratase | 0.65 |
| P07897-2 | Isoform 2 of Aggrecan core protein | 0.65 |
| P23928 | Alpha-crystallin B chain | 0.64 |
| P00564 | Creatine kinase M-type | 0.63 |
| P14141 | carbonic anhydrase 3 | 0.62 |
| P07323 | Gamma-enolase | 0.61 |
| P04466 | Myosin regulatory light chain 2, skeletal muscle isoform | 0.59 |
| P56571 | ES1 protein homolog, mitochondrial | 0.59 |
| Q9QZ76 | Myoglobin | 0.58 |
| P02600-1 | Myosin light chain 1/3, skeletal muscle isoform | 0.53 |
| P16409 | myosin light chain 3 | 0.53 |
| Q01717 | Thyrotropin-releasing hormone receptor | 0.43 |
| Q9EPH1 | Alpha-1B-glycoprotein | 0.37 |

**Table S3: Identified proteins showing ≥1.5 fold change between AIA and TXR200 study groups.**

| **Accession no** | **Protein name** | **Ratio (AIA/TXR200)** |
| --- | --- | --- |
| P19132 | Ferritin heavy chain | 1.89 |
| P13084 | Nucleophosmin | 1.75 |
| P01048 | T-kininogen 1 | 1.71 |
| Q62930 | complement component C9 | 1.70 |
| P08932 | T-kininogen 2 | 1.69 |
| P48199 | C-reactive protein | 1.64 |
| Q5I0G4 | Glycine--tRNA ligase | 1.61 |
| P17475 | Alpha-1-antiproteinase | 1.60 |
| P50116 | Protein S100-A9 | 1.53 |
| P23928 | Alpha-crystallin B chain | 0.66 |
| Q9EPH1 | Alpha-1B-glycoprotein | 0.60 |

**Table S4: List of significantly differentially deregulated proteins between AIA and TXR200.**

| **Accession no** | **Protein Name** | **p-value (t-test; AIA/TXR200)** |
| --- | --- | --- |
| P02454 | Collagen alpha-1(I) chain | 0.0003 |
| Q6P6Q2 | keratin, type II cytoskeletal 5 | 0.004 |
| Q62930 | complement component C9 | 0.007 |
| O88767 | protein/nucleic acid deglycase DJ-1 | 0.008 |
| P17475 | Alpha-1-antiproteinase | 0.008 |
| P23965 | Enoyl-CoA delta isomerase 1, mitochondrial | 0.010 |
| O35952-1 | Hydroxyacylglutathione hydrolase, mitochondrial | 0.015 |
| Q63416 | Inter-alpha-trypsin inhibitor heavy chain H3 | 0.01 |
| P00507 | Aspartate aminotransferase, mitochondrial | 0.01 |
| P48199 | C-reactive protein | 0.01 |
| B2RYG6 | Ubiquitin thioesterase otub1 | 0.02 |
| P50878 | 60S ribosomal protein L4 | 0.02 |
| Q811A3-2 | Isoform 2 of Procollagen-lysine,2-oxoglutarate 5-dioxygenase 2 | 0.02 |
| Q5XIM9 | T-complex protein 1 subunit beta | 0.02 |
| O35763 | Moesin | 0.03 |
| P04762 | catalase | 0.03 |
| P11884 | Aldehyde dehydrogenase, mitochondrial | 0.03 |
| Q9EPH1 | Alpha-1B-glycoprotein | 0.03 |
| P07150 | annexin A1 | 0.03 |
| Q5U300 | Ubiquitin-like modifier-activating enzyme 1 | 0.03 |
| P04642 | L-lactate dehydrogenase A chain | 0.03 |
| P13437 | 3-ketoacyl-CoA thiolase, mitochondrial | 0.03 |
| Q5XI78 | 2-oxoglutarate dehydrogenase, mitochondrial | 0.04 |
| O35567 | bifunctional purine biosynthesis protein purH | 0.04 |
| P51886 | Lumican | 0.04 |
| P20767 | Ig lambda-2 chain C region | 0.04 |
| Q63617 | Hypoxia up-regulated protein 1 | 0.049 |

**Table S5: Identified proteins showing ≥1.5 fold between TXR and DS study groups with respect to Healthy.**

| **Accession** | **Protein name** | **Ratio (TXR/Healthy)** | **Ratio (DS/Healthy)** |
| --- | --- | --- | --- |
| P02764 | Alpha-1-acid glycoprotein | 2.51 | 3.8 |
| P02650 | Apolipoprotein E | 2.24 | 2.23 |
| P10960 | Prosaposin | 2.19 | 4.17 |
| P63088-2 | Isoform Gamma-2 of Serine/threonine-protein phosphatase PP1-gamma catalytic subunit | 2.02 | 1.95 |
| P24268 | Cathepsin D | 1.98 | 2.81 |
| P20059 | Hemopexin | 1.94 | 2.14 |
| P06302 | Prothymosin alpha | 1.91 | 2.7 |
| Q62829 | Serine/threonine-protein kinase PAK 3 | 1.78 | 2.7 |
| P07895 | Superoxide dismutase [Mn], mitochondrial | 1.77 | 3.4 |
| P80067 | Dipeptidyl peptidase 1 | 1.63 | 2.26 |
| P34058 | Heat shock protein HSP 90-beta | 1.61 | 1.66 |
| P61983 | 14-3-3 protein gamma | 1.60 | 1.64 |
| Q63797 | Proteasome activator complex subunit 1 | 1.59 | 1.94 |
| Q6P686 | osteoclast-stimulating factor 1 | 1.58 | 2.26 |
| Q6P6Q2 | keratin, type II cytoskeletal 5 | 1.57 | 1.86 |
| P01041 | Cystatin-B | 1.56 | 2.05 |
| D3ZHA0 | Filamin-C | 1.56 | 1.6 |
| Q6IFV3 | Keratin, type I cytoskeletal 15 | 1.55 | 2.007 |
| P20759 | Ig gamma-1 chain C region | 1.55 | 1.54 |
| P05942 | Protein S100-A4 | 1.51 | 1.41 |
| Q62930 | complement component C9 | 1.5 | 1.77 |
| P11598 | Protein disulfide-isomerase A3 | 1.5 | 2.1 |
| P18420 | Proteasome subunit alpha type-1 | 1.5 | 1.84 |
| P05065 | fructose-bisphosphate aldolase A | 1.5 | 1.49 |
| P11980-1 | Pyruvate kinase PKM | 0.64 | 0.6 |
| Q9EPH1 | Alpha-1B-glycoprotein | 0.61 | 0.39 |
| P56571 | ES1 protein homolog, mitochondrial | 0.56 | 0.58 |
| Q01717 | Thyrotropin-releasing hormone receptor | 0.49 | 0.38 |

| **S. No.** | **Gene ID** | **Uniprot accession number** | **Protein** | **Interacting molecule** | **Binding Affinity**  **(Kcal / mol.)** | **Interacting Residues** |
| --- | --- | --- | --- | --- | --- | --- |
| 1 | C9 | Q62930 | Complement component 9 | Troxerutin | -8.9 | ALA86, ALA201, GLY193, GLU77, PRO85 |
| 2 | Lta4h | P30349 | Leukotriene A4 hydrolase |  | -8.5 | TYR267, ALA378, SER379, SER380, TYR383, ARG537, ALA504, ASN351, GLU348, PRO569, ASP606 |
| 3 | Pfkp | P47860 | Phosphofructokinase, platelet |  | -8.4 | ARG(348, 385, 725), SER386, ASP(182, 726), LEU727, THR(181, 393), GLY(180, 389), LYS688, TYR223 |
| 4 | Pdia3 | P11598 | Protein disulfide isomerase family A, member 3 |  | -8.0 | ASN181, GLY123, SER126, TYR182, ARG183, LYS(130, 152), ASP250 |
| 5 | Afm | P36953 | Afamin |  | -7.9 | PHE(135, 139,158), ASP542, TYR162, ASN132,170, THR210, ARG169 |
| 6 | Dhfr | Q920D2 | Dihydrofolate reductase |  | -7.5 | ILE17, GLY21, LEU76, ARG55, SER60, LYS56 |
| 7 | Aldh2 | P11884 | Aldehyde dehydrogenase 2 family |  | -7.3 | ALA100, SER101, HIS175, ARG(103,174), GLY160, THR173 |
| 8 | Vps29 | B2RZ78 | VPS29 retromer complex component |  | -7.2 | ILE91, LEU102, SER113, ILE85 |
| 9 | Cat | P04762 | Catalase |  | -7.1 | MET339, ASP140, ARG382, ASN142, GLN414 |
| 10 | Ppp1cc | P63088 | Protein phosphatase 1 catalytic subunit gamma |  | -6.7 | CYS273, ARG96, LYS98, GLU300, ALA299, ASN271, GLY274, ASP277 |
| 11 | Gyg1 | O08730 | Glycogenin 1 |  | -6.7 | LEU183, VAL186, PHE(185, 204), ALA200, TYR197 |
| 12 | Trhr | Q01717 | Thyrotropin releasing hormone receptor |  | -6.6 | THR60, ASN61, PHE119, ARG123, ALA126, ILE127, PRO130, VAL264, MET267, SER324, GLN325, LYS326 |
| 13 | Lhpp | Q5I0D5 | Phospholysine phosphohistidine inorganic pyrophosphate phosphatase |  | -6.4 | ALA(86, 201), GLY193, GLU77, PRO85 |
| 14 | Rac1 | Q6RUV5 | Ras-related C3 botulinum toxin substrate 1 |  | -6.3 | ALA13, GLY15, THR17, CYS18, TYR32, SER83 |
| 15 | SerpinA6 | P31211 | Serpin family A member 6 |  | -5.9 | LEU312, THR(279, 313), ASP(160, 278), LYS34, VAL155 |

**Table S6.** *In silico* key regulators derived using the community finding algorithm from all identified proteins and their binding affinity towards TXR.
